# *Mycobacterium tuberculosis* employs atypical and different classes of B_12_ switches to control separate operons

**DOI:** 10.1101/2023.04.25.538288

**Authors:** Terry Kipkorir, Peter Polgar, Declan Barker, Alexandre D’Halluin, Zaynah Patel, Kristine B. Arnvig

**Affiliations:** Institute for Structural and Molecular Biology, University College London, Gower Street, WC1E 6BT London, United Kingdom; Department of Infection Biology, The London School of Hygiene and Tropical Medicine, Keppel Street, WC1E 7HT London, United Kingdom; Institut de Biologie Physico-Chimique, Université de Paris, 75005 Paris, France

## Abstract

Vitamin B_12_ (B_12_), an essential cofactor in all domains of life, is produced *de novo* by only a small subset of prokaryotes, but B_12_-sensing riboswitches are some of the most widely distributed riboswitches in bacteria. *Mycobacterium tuberculosis*, the causative agent of the ongoing tuberculosis pandemic, encodes two distinct vitamin B_12_ riboswitches. One controls the expression of *metE*, encoding a B_12_-independent methionine synthase, while the other is located upstream of *ppe2,* a PE/PPE family gene whose function is still unresolved. Here, we analyse ligand sensing, secondary structure architecture, and gene expression control mechanisms of these two riboswitches. Our results provide the first evidence of direct ligand binding by *metE* and *ppe2* riboswitches and show that the two switches exhibit different preferences for natural isoforms of B_12_, use distinct regulatory and structural elements, and act as translational OFF switches. Based on our results, we propose that the *ppe2* switch represents a new Class IIc of B_12_-sensing riboswitches. Moreover, we have identified small translated open reading frames (uORFs) upstream of both *metE* and *ppe2*, which modulate the expression of the respective downstream genes in opposite directions. Translation of the *metE* riboswitch uORF suppresses MetE expression, while translation of the uORF in the *ppe2* switch is essential for PPE2 expression via the synthesis of a uORF-PPE2 fusion protein. In summary, our findings reveal an unexpected diversity and complexity of B_12_-dependent *cis*-regulation in *M. tuberculosis*, with potential implications for host-pathogen interactions.

## Introduction

RNA leaders preceding coding sequences in mRNAs have gained interest as hubs for gene expression control (1–4). These include riboswitches, which are highly structured *cis*-regulatory RNAs that sense and bind specific metabolites such as enzyme cofactors, amino acids, or nucleotides, to affect the expression of genes under their control (5–7). Typically, the regulated genes have a direct relationship with the corresponding riboswitch ligand, whereby the encoded gene product is involved in the *de novo* biosynthesis of the ligand or its transport (8, 9). Riboswitches regulate gene expression via the interaction between two RNA domains: the aptamer and the expression platform. The aptamer is highly conserved and forms a unique three-dimensional structure for ligand-binding. The expression platform executes gene regulation by adopting mutually exclusive secondary structures depending on the ligand binding status of the aptamer (3, 4). Although the mechanisms of individual riboswitches vary, the gene expression outcome is either permissive (“ON” switch) or non-permissive (“OFF” switch), primarily resulting from changes in transcription termination or translation initiation or both (10–12). Termination of transcription can be either intrinsic or Rho-dependent. Intrinsic transcription termination requires the formation of a stable hairpin, which in the context of a U-rich tail leads to dissociation of the elongation complex (13, 14). In the Rho-dependent mechanism, the ATP-dependent RNA helicase Rho binds to exposed C-rich sequences on RNA (known as Rho utilisation (*RUT*) sites), and translocates along the transcript to disrupt the elongation complex further downstream (15–17). Riboswitch-mediated translational control typically involves ligand-dependent occlusion of the Shine-Dalgarno sequence (SD) by a complementary anti-SD (αSD) sequence, which, in turn, may be sequestered by an anti-anti-SD (ααSD) sequence (12). The absence of translation may in turn expose *RUT* sites on the mRNA, facilitating Rho-dependent transcription termination (16, 18–20). Intrinsic terminators are rare in mycobacteria and therefore, mycobacterial riboswitches are presumably mostly translational and/or Rho-dependent (4, 21).

Numerous riboswitch families regulating a wide range of cellular processes have been discovered in the last two decades (22–31). One of the most widespread families of riboswitches bind coenzyme B_12_ (cobalamin), regulating genes involved in the biosynthesis, transport or utilisation of this cofactor (31–34). B_12_ is a complex organometallic compound whose structure is based on a corrin ring containing a central cobalt ion coordinated by lower (α-axial) and upper (β-axial) ligands (**Fig. 1**) (35). The α-axial ligand is typically dimethylbenzimidazole (DMB)(36), while the β-axial ligand can be one of several eponymous functional groups (“R” group), among which adenosyl (Ado-), methyl (Me-), hydroxy (Hy-), and cyano (CN-) groups are the most common (**Fig. 1**) (35). Adenosylcobalamin (AdoB_12_), methylcobalamin (MeB_12_), and hydroxocobalamin (HyB_12_) are naturally occurring B_12_ isoforms, whereas cyanocobalamin (CNB_12_) is predominantly produced in large scale as a derivative of the natural B_12_ isoforms in *Pseudomonas denitrificans*, *Propionibacterium shermanii*, or *Sinorhizobium meliloti* (37).

**Figure 1.**
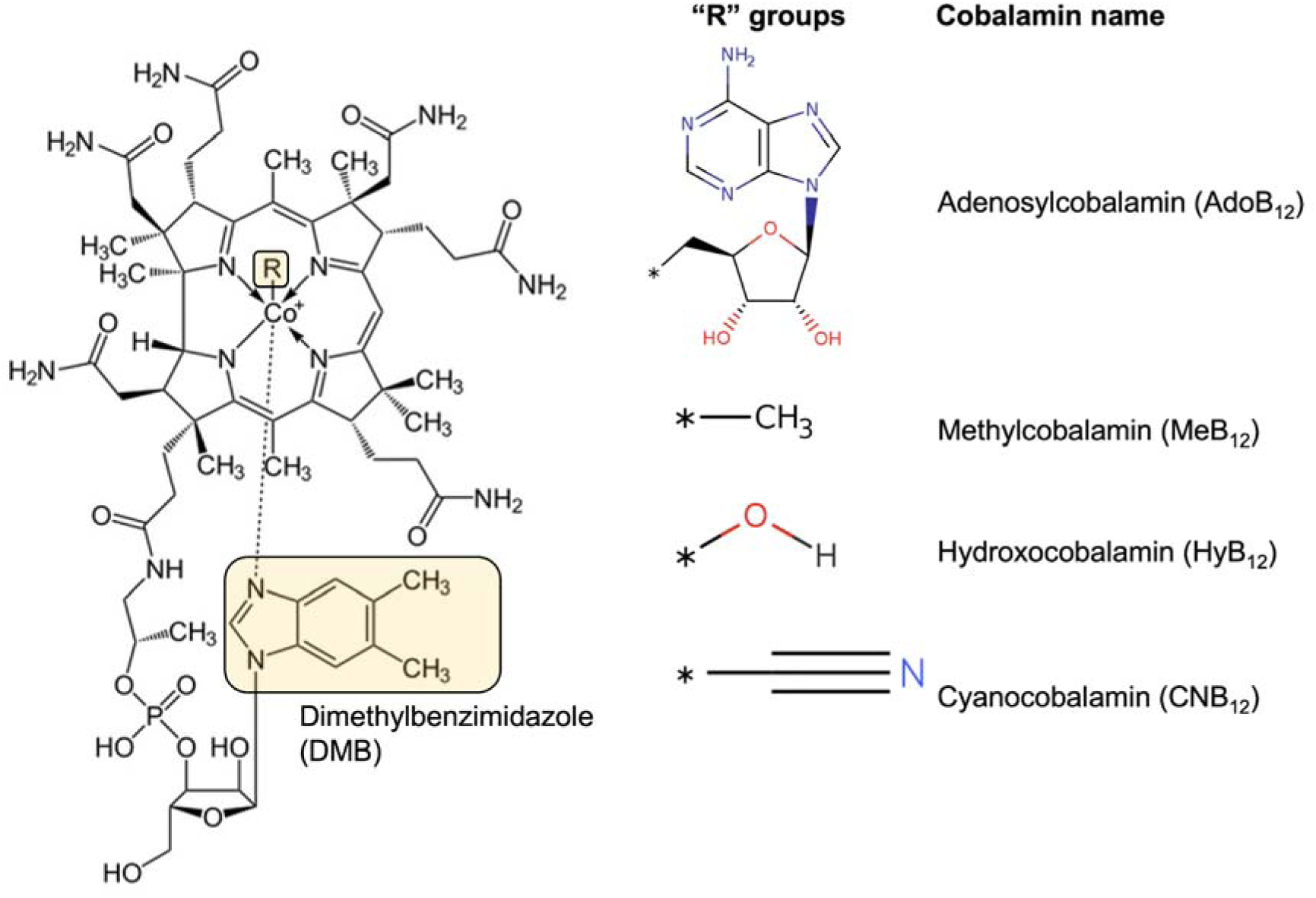
Chemical structure of cobalamin. The different functional groups (R groups) that occupy the β-axial positions are shown to the right of the structure and the α-axial base, dimethylbenzimidazole (DMB), is highlighted on the structure.

*Mycobacterium tuberculosis*, the aetiological agent of tuberculosis (38, 39), lacks the ability to synthesise B_12_ (40, 41) due to the deletion of *cobF*, encoding a precorrin-6A synthase involved in the *de novo* B_12_ biosynthesis pathway, which is otherwise intact (42–44). Still, *M. tuberculosis* encodes several B_12_-dependent enzymes, suggesting that the pathogen utilises host-derived B_12_ whenever available (45, 46), although these enzymes are found alongside alternative B_12_-independent counterparts for the same metabolic processes. The conversion of ribonucleotides to deoxyribonucleotides may be catalysed by the B_12_-independent ribonucleotide reductase NrdEF or by the AdoB_12_-dependent enzyme, NrdZ (45, 46). Similarly, the degradation of propionate, a toxic by-product of odd-chain fatty acid catabolism may be accomplished via the B_12_-independent methyl citrate cycle or by the methyl malonate pathway, which includes the AdoB_12_-dependent methylmalonyl-CoA mutase MutAB (45, 46). Finally, the biosynthesis of methionine from homocysteine may be catalysed by the B_12_-independent methionine synthase, MetE or by the MeB_12_-dependent enzyme, MetH (45, 46). Methionine synthesis and propionate degradation are both B_12_-dependent pathways in humans (47).

B_12_ has been shown to regulate expression of *metE* via B_12_-sensing riboswitches in *M. tuberculosis* and in the saprophytic mycobacterial model, *Mycobacterium smegmatis*, by suppressing *metE* mRNA levels (48, 49). A second, homologous B_12_ riboswitch occurs in *M. tuberculosis* upstream of a potential three-gene operon comprising *ppe2*, *cobQ* and *cobU*. PPE2 is a member of the large, Mycobacterium-specific PE/PPE protein family associated with host-pathogen interactions and virulence (50–52); specifically, PPE2 is suspected to be involved in cobalt transport, whereas CobQ and CobU are relics of the disrupted B_12_ biosynthesis pathway (53).

Biochemical and structural studies of B_12_ riboswitches in different bacteria have provided detailed insights into conserved features of this element (54–57). B_12_ riboswitches share a conserved ligand-binding core formed of a central four-way junction and peripheral elements egressing from this junction (31, 58). The interplay between aptamer and expression platform involves an interdomain “kissing loop”, whose formation is thought to be strictly ligand-driven (57, 58). Despite the conservation of these features, B_12_ riboswitches exhibit differential selectivity to the B_12_ isoforms, arising from sequence and structural variations in the peripheral elements (57, 58). Consequently, B_12_ riboswitches have been broadly classified into three distinct classes (Class I, IIa & IIb). Class I and IIb riboswitches selectively bind AdoB_12_, whereas Class IIa riboswitches show preferential binding to the slightly smaller MeB_12_ and HyB_12_. Other B_12_ riboswitches displaying promiscuous binding to a broad range of corrinoids have also been discovered (53). Given that only a small fraction of known riboswitches have been characterised and thousands more are probably still undiscovered (33, 59), it is reasonably expected that novel regulatory mechanisms of B_12_ riboswitches will continue to be uncovered across a wide range of species.

Recently, it has been proposed that codons and/or peptides arising from the translation of so-called upstream open reading frames (uORFs) occurring within gene leaders can either positively or negatively impact the expression level of the downstream ORF (21, 60–62). For example, the presence of translating ribosomes on uORFs in mRNA 5’ regions can modulate the expression of the downstream gene by inducing or suppressing premature transcription termination, or by interfering with translation initiation of the downstream ORF (63). In addition, if the uORF either overlaps with or occurs in the same reading frame of the downstream ORF, some form of translation coupling between the ORFs may occur (64, 65). How uORFs embedded within riboswitches might affect their function has not yet been addressed.

In the current study, we analyse ligand binding, riboswitch architecture, and control mechanisms of the *metE* and *ppe2* riboswitches from *M. tuberculosis*. Our results provide the first evidence of direct binding of B_12_ to these elements and the functional validation of the *ppe2* riboswitch as an “OFF switch.” We found that the two riboswitches exhibit differential binding of AdoB_12_, MeB_12_, HyB_12_ and CNB_12_, and involve distinct features in their expression platforms to execute B_12_-dependent control. On this basis, we propose that the *ppe2* switch represents a new (Class IIc) switch. Moreover, we show that the translation of uORFs in the leaders of *metE* and *ppe2* alters the expression of their respective downstream annotated coding regions. Interestingly, translation of the *metE* uORF suppresses MetE expression whereas translation of the *ppe2* uORF is essential for PPE2 expression. In the latter case, the uORF stop codon is suppressed, resulting in a frameshifted uORF-PPE2 fusion protein. These findings expand our current understanding of B_12_ riboswitches by exposing their variations and complexity within a single organism.

## Results

### Premature transcription termination in metE and ppe2 leaders

We have mapped multiple premature transcription termination sites (TTS) associated with RNA leaders including those of *metE* and *ppe2* in *M. tuberculosis* cultures grown in standard conditions (21). The TTS patterns within these two leaders indicated multiple TTS thoughout the *metE* leader, compared to two, closely spaced TTS in the *ppe2* leader, suggesting differences in the regulation of the switches (**Fig. 2A** **& B**) (21). To explore how B_12_ might affect growth and *metE* and *ppe2* transcription, we grew cultures of *M. tuberculosis* to OD_600_∼0.6 before adding 10 μg/mL AdoB_12_; notably, this did not affect the growth rate (**Supplementary Fig. 1**). RNA was isolated before and 1 hour after the addition of AdoB_12_, and analysed by quantitative RT-PCR (qRT-PCR) targeting the leader and the coding regions of *metE* and *ppe2* (**Fig. 2A** **& B**). The results suggested that addition of AdoB_12_ led to a 19-fold decrease in the level of *metE* leader RNA and 60-fold decrease in *metE* coding RNA (**Fig. 2C**). The changes in *ppe2* transcript levels were more modest by comparison, with *ppe2* leader RNA decreasing ∼2-fold and coding RNA reducing ∼2.5-fold following AdoB_12_ addition (**Fig. 2D**).

**Figure 2.**
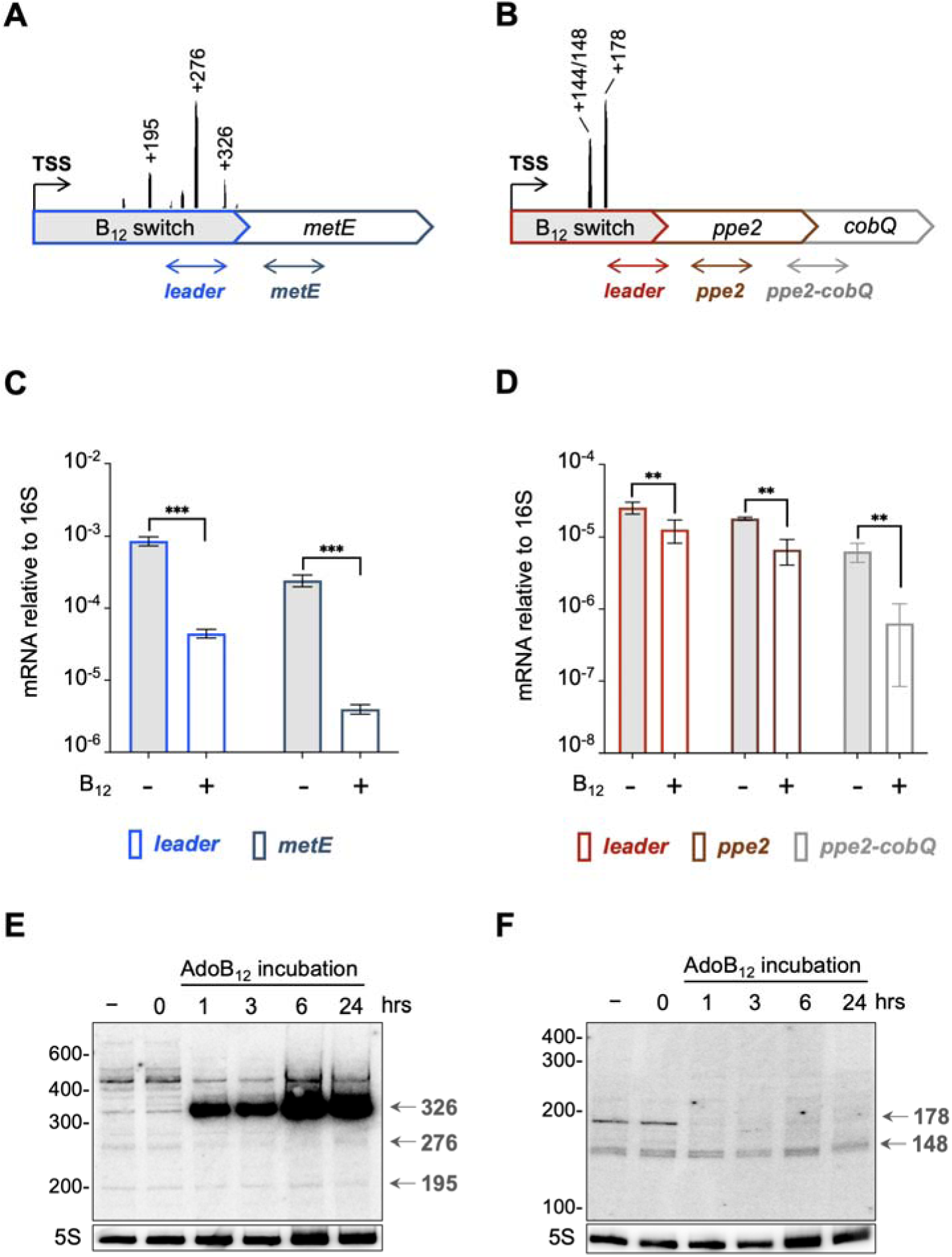
B_12_-dependent changes in *metE* and *ppe2* transcripts. **A-B.** Schematic of *metE* and *ppe2* operons; full-length of leaders are 365 nt and 292 nt, respectively (not drawn to scale with coding regions). Dominant transcription termination sites (TTS) and their relative coverage according to (21) indicated. Approximate locations of qRT-PCR amplicons for *metE*/*ppe2* leaders, coding regions and *ppe2-cobQ* junction are indicated as double-ended arrows; exact locations are listed in methods. **C-D.** qRT-PCR of amplicons indicated in A and B before and 1 h post AdoB_12_ addition; note different scales. Data represents mean ± standard deviation of at least three biological replicates, *p*-values *t*-test: ** *p*<0.05; *** *p*<0.001. **E-F.** Northern blots showing transcript changes in *metE* (**E**) and *ppe2* (**F**) leaders before and after AdoB_12_ incubation with probes hybridising to 5’ end of mRNA. Arrows indicate signals corresponding to TTS peaks indicated in A and B.

The *cobQU* genes downstream of *ppe2* are associated with cobalamin synthesis in other species (35, 53), making them likely targets of B_12_-dependent control. To determine if *cobQ* was co-transcribed with *ppe2* and thus potentially regulated by the B_12_ switch, we also performed qRT-PCR across the *ppe2-cobQ* junction. The results indicated that *ppe2* and *cobQ* are indeed co-transcribed and that RNA levels decrease 10-fold after AdoB_12_ addition, suggesting that *cobQ* is also regulated by the riboswitch (**Fig. 2B** **& D).** RT-PCR did not amplify across the *cobQ-cobU* junction, implying that *cobU* is not part of this operon (**Supplementary Fig. 1**). In summary, the results suggest transcriptional polarity in both *loci*, albeit much less in *ppe2* than in *metE*.

To validate the notion of premature termination, we performed northern blotting of RNA at times 0, 1, 3, 6 and 24 hours post B_12_-addition using a probe that hybridised to the 5’ end of each leader. The transcript pattern before B_12_-addition reflected the TTS mapping with multiple signals for *metE* and only a few for *ppe2*; moreover, some of the signals on the blots corresponded to the dominant TTS signals indicated in panels **A** and **B** (**Fig. 2E** **& F**, grey arrows). The addition of AdoB_12_ led to one primary but seemingly opposite B_12_-dependent change within each leader: the 326-nt signal within the *metE* leader increased substantially upon AdoB_12_ addition, suggesting premature termination of transcription (**Fig. 2E**), while the 178-nt signal within the *ppe2* leader disappeared (**Fig. 2F**). These results further support the notion of B_12_-dependent termination of transcription in the *metE* leader. The *ppe2* results are somewhat ambiguous but nevertheless corroborate substantial differences between the two riboswitches.

### The metE and ppe2 aptamers display variable selectivity for B_12_ isoforms

To ascertain direct interactions between B_12_ and the *metE* and *ppe2* aptamers and to determine the ligand binding properties of each, we generated transcripts for inline probing analysis (66) by *in vitro* transcription. Both transcripts covered the region between the TSS and a few bases downstream of their respective most distal leader TTS. Thus, the *metE* riboswitch transcript was 345 nucleotides long, whereas the *ppe2* transcript was 191 nucleotides in size. First, we analysed the interactions between the two riboswitches and four common B_12_ isoforms: AdoB_12_, MeB_12_, HyB_12_ & CNB_12_. Inline probing reactions contained at least a 4-log excess concentration of ligand (1 mM) over that of RNA. In the case of *metE*, AdoB_12_ induced the strongest modulation signals, while MeB_12_, HyB_12_ or CNB_12_ resulted in little to no change (**Supplementary Fig. 2**). In contrast, all four B_12_ isoforms resulted in similar cleavage patterns and signal intensities in the *ppe2* transcript with the possible exception of C48-C50, where only AdoB_12_ caused a reduction in the cleavage signal (**Supplementary Fig. 2**). In summary, our results indicate that the regions flanked by the TSS and the distal TTS are sufficient for ligand binding in both riboswitches and while the *metE* riboswitch is selective for AdoB_12_, the *ppe2* riboswitch seems able to accommodate all four B_12_ isoforms equally well.

### B_12_-binding leads to occlusion of the metE translation initiation region

To interrogate the ligand sensitivity of the *metE* riboswitch, we performed inline probing using a range of AdoB_12_ concentrations from 1 nM to 2 mM (**Fig. 3A**), which suggested an approximate dissociation constant (K_d_) <60 μM (range = 5 μM – 57 μM) (**Supplementary Fig. 3**). The inline probing data were used to apply constraints to a predicted structure of the ligand-bound switch on the RNAstructure web server (67). The resulting structure indicated that the aptamer domain of the *metE* riboswitch is contained within the first 220 nucleotides of the leader largely in agreement with the prediction in Rfam (68). Hence, the region downstream of this position was considered part of the expression platform. Several regions of hypercleavage, including G240/C245 & C255/G265, occur in the proposed expression platform in the presence of AdoB_12_ (**Fig. 3A**). These positions are paired in the predicted unbound structure (Apo-form) but adopt single-stranded conformations in the ligand-bound state (**Fig. 3B** **& C**). An additional structure not reported in Rfam (68) was predicted at positions 1-14, comprising a short hairpin (P0) that is highly conserved in mycobacteria (**Fig. 3C**; **Supplementary Fig. 4**).

**Figure 3.**
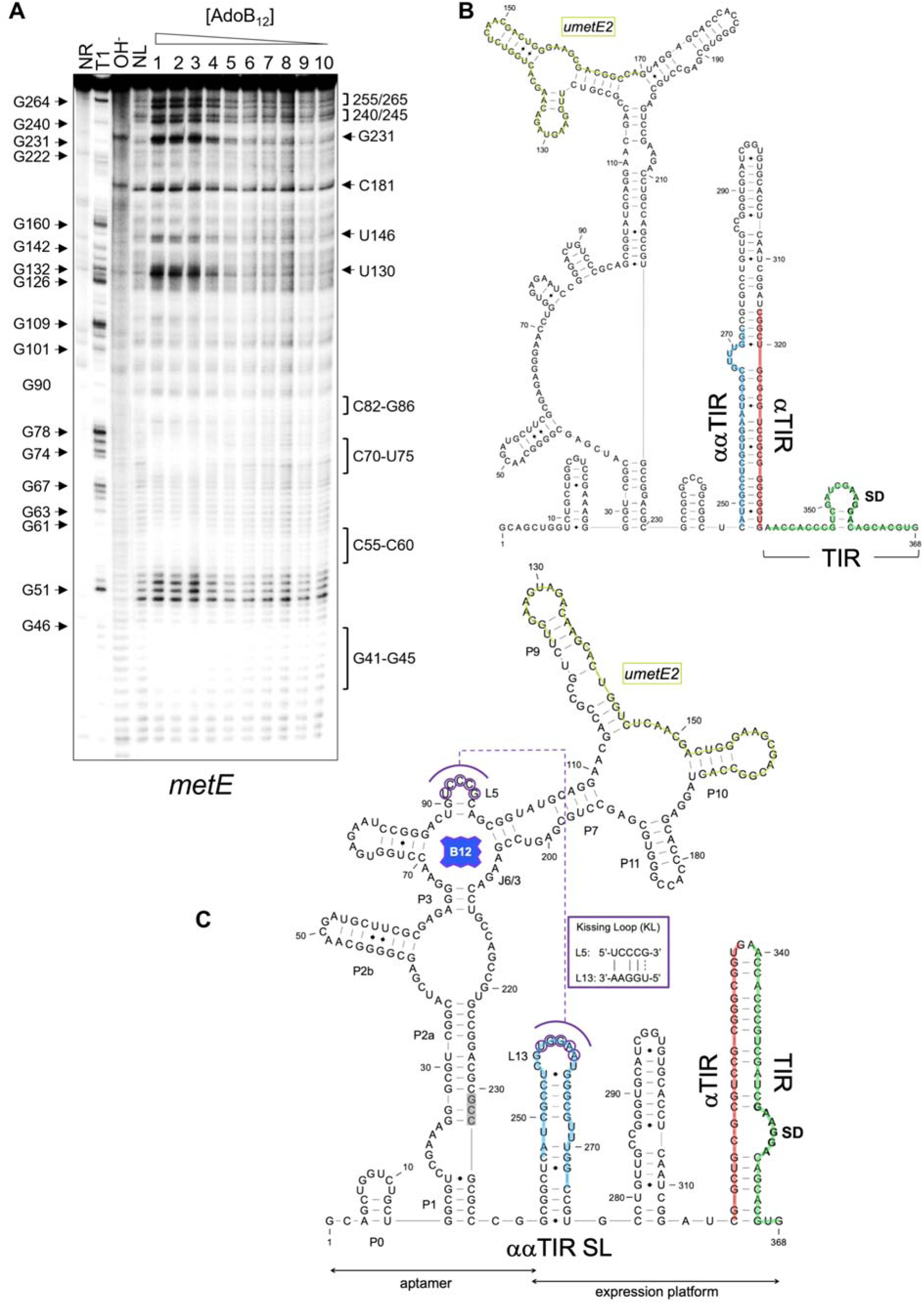
Inline probing uncovers unusual features of *metE* riboswitch. **A.** Cleavage pattern of the *metE* riboswitch RNA over a concentration gradient of AdoB_12_ (lanes, 1: 2 mM; 2: 1 mM; 3: 0.5 mM; 4: 0.1 mM; 6: 10 uM; 6: 1 uM; 7: 0.1 uM; 8: 10 nM; 9: 1 nM; 10: 0.1 nM. T1 – RNAse T1; OH^-^ – alkaline digest; NL – no-ligand control; NR – no-reaction. Strongly modified positions are indicated on the right side of the gel, (cropped where there was no difference ± ligand). T1-derived G positions indicated on the left. **B.** Predicted structure of the *metE* switch without ligand. The translation initiation region (TIR) including the SD sequence is highlighted in green, while the αTIR and ααTIR sequences are highlighted in red and blue, respectively. uMetE2 is described in the text. **C.** Predicted structure of AdoB_12_-bound *metE* switch. Paired regions in the aptamer are labelled sequentially (P0-P11). The conserved “B_12_ box” (G199-C215) matches the consensus for this element (68). Bases of L5 that could potentially form a kissing loop (KL) with L13 are highlighted in purple and the base pairs between the loops are shown in the inset. The region of hypercleavage at G231-C233 is highlighted in grey. Other coloured features are the same as in panel B.

According to the predicted structure, the B_12_ binding pocket is enclosed in a four-way junction formed by P3-P6. The conserved “B_12_ box” (31) in this switch stretches from G199 to C215, and is followed immediately downstream by a hypercleaved positions at G231-C233 (**Fig. 3A** **& C**). Issuing from the B_12_ pocket are the peripheral elements comprising a bipartite P2 arm and a large P6 extension featuring a second four-way junction formed by P7, P8, P10 & P11 (**Fig. 3C**). Moreover, a stereotypical “kissing loop” (KL) is formed by pseudoknot interactions between conserved CCC nucleotides in L5 and a variable partner loop located in the expression platform. To validate the assignment of L5, we substituted the CCC with AAA, which led to a loss of regulation, suggesting that this was indeed L5 and part of the KL (data not shown).

The typical expression platform of B_12_ riboswitches is translational (31), and we found no evidence of an intrinsic terminator within the *metE* switch. Comparing the ligand-bound and Apo-structures in **Fig. 3B and C** we identified a potential expression platform, in which a broad translation initiation region (TIR) including 25 nucleotides upstream of the MetE start codon (GUG) and a likely SD were effectively occluded by a complementary α-TIR stem in the ligand-bound structure (**Fig. 3C**). In the Apo-form, the same α-TIR was fully sequestered by extensively pairing with an αα-TIR, thereby unmasking the TIR (**Fig. 3B**). Finally, the αα-TIR formed a hairpin in which the apical loop (GUGGA) presented an ideal pairing partner for L5, thereby linking the KL directly to elements of the expression platform (**Fig. 4C**). This structure and the ligand preference for AdoB_12_ suggested that the *metE* switch is a Class I switch.

**Figure 4.**
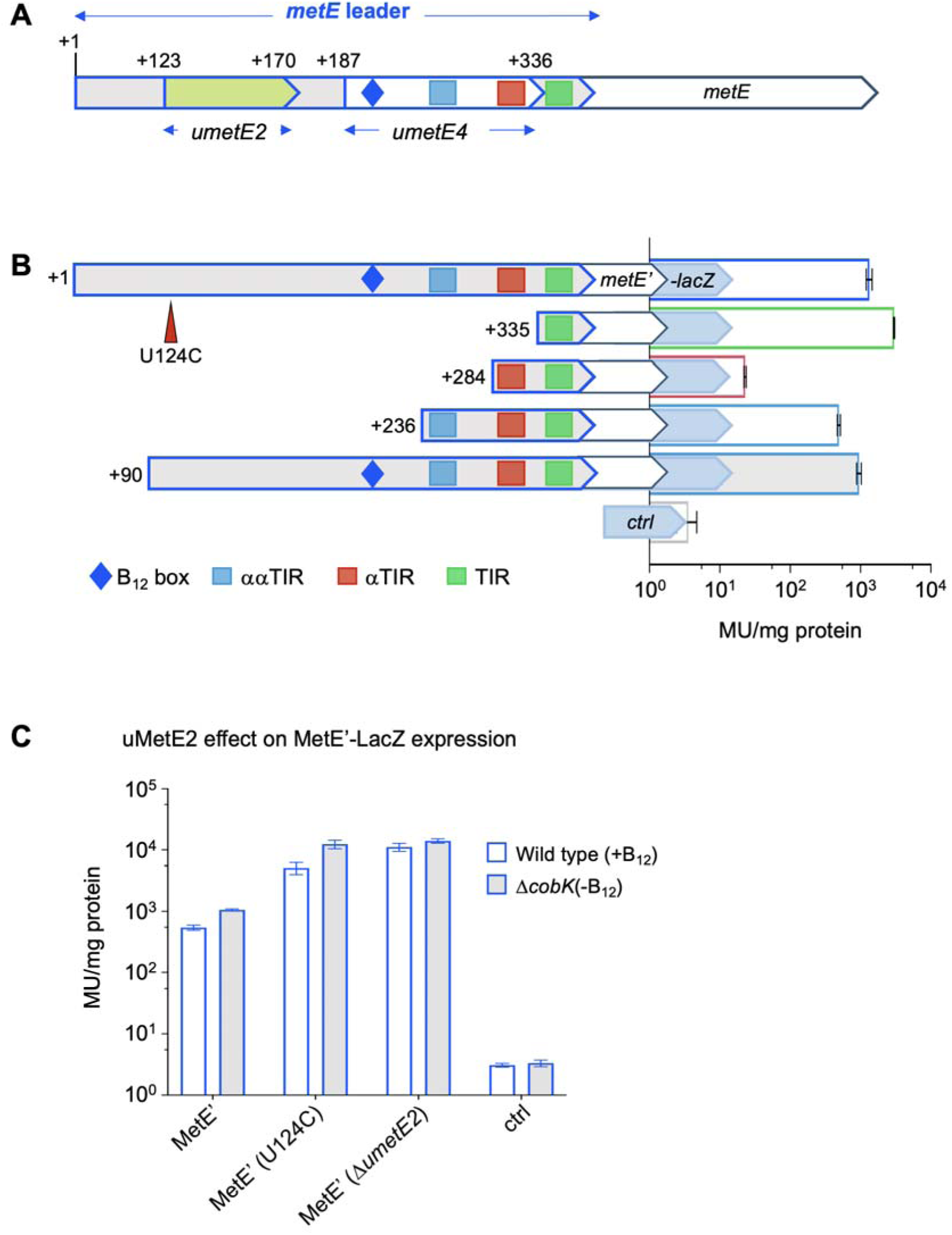
Translation of uMetE modifies MetE expression. **A** Outline of the *metE* leader including control elements (TIR, αTIR and ααTIR) identified in Fig. 3 and the potentially translated leader-encoded peptides, uMetE2 and uMetE4. **B.** Outline of reporter constructs for the validation of the translational expression platform using gradual 5’ extensions; numbers indicate 5’ ends of each construct. β-gal activities are in Miller Units (MU)/mg protein. ctrl: no SD. **C.** Assessing the role of uMetE2 translation. The expression of MetE’-LacZ was measured in the context of wild type *umetE2*, non-translated *umetE2* (U124C) or Δ*umetE2* (Δ124-180) all expressed in wild type *M. smegmatis* (B_12_ producing) or Δ*cobK* (B_12_ deficient). Data represents mean ± standard deviation of at least three biological replicates.

To validate these elements, we made a series of reporter constructs, in which the second codon of *metE* was fused in frame to *lacZ* (*metE’-lacZ*). The upstream edge consisted of gradual 5’ extensions of the *metE* leader to include the predicted TIR, αTIR and the ααTIR, respectively (**Fig. 4**). Predicted structures of these partial leader-constructs are shown in **Supplementary Fig. 5**. To avoid B_12_-dependent folding affecting β-galactosidase (β-gal) activities, all constructs were transformed into a *M. smegmatis* Δ*cobK* mutant unable to synthesise B_12_ (49). The results indicated that the level of MetE expression in the minimal construct (TIR only) was much higher than that of the full-length *metE* leader, suggesting the presence of inhibitory sequences upstream of the basic TIR (**Fig. 4B**). Extending the construct to include the predicted αTIR sequence reduced MetE’-LacZ expression substantially, supporting the notion of translation inhibition via TIR occlusion. A further extension to include the ααTIR sequence partially reversed this phenotype, further supporting the predicted structure-function relationship (**Fig. 3** **&** **Fig. 4**). Finally, an extension as far back as +90 restored MetE’-LacZ expression level to nearly that of the full-length switch. These findings suggest that the *metE* riboswitch is an “OFF” switch that employs TIR occlusion (*i.e*., a translational expression platform) via the suggested elements for gene expression control. The formation of the KL would directly block the proposed ααTIR and reinforce the αTIR-TIR pairing. However, as previously noted, we also observed a B_12_-enhanced TTS within the *metE* leader and a significant reduction in *metE* RNA levels, suggesting that the translational expression platform is augmented by Rho-dependent termination of transcription and/or rapid mRNA degradation following reduced translation.

### A short translated ORF within the metE aptamer suppresses MetE expression

*M. tuberculosis* encodes numerous short, translated ORFs (uORFs) upstream of annotated genes including genes controlled by riboswitches (21, 69). The *metE* riboswitch encodes four potentially translated uORFs (*umetE1*-*4*) based on credible SD sequences and associated start codons at +88 (CUG), +123 (UUG), +140 (CUG), & +187 (GUG), relative to the TSS (**Fig. 3**). Translation within a riboswitch aptamer will likely impact its folding and hence, the function of the switch. To investigate whether the presence of *metE* uORFs did affect riboswitch function, we investigated these in more detail. As translation in mycobacteria rarely initiates with CUG start codons (21, 69–71), we pursued only *umetE*2 (+123 to +170) and the larger *umetE4* (+187 to +336) as potentially translated uORFs (**Fig. 3** **&** **Fig. 4**). In-frame reporter fusions, covering the region upstream of the SD to 8 codons of uMetE2 (uMetE2’-LacZ) or 5 codons of uMetE4 (uMetE4’-LacZ), respectively, were expressed in wild type (B_12_ proficient) *M. smegmatis*. The resulting β-gal assays indicated low levels of uMetE2’-LacZ expression (27 ± 1.4 MU/mg protein) compared to the much more highly expressed MetE’-LacZ (544 ± 52.1 MU/mg protein), whereas the expression of uMetE4’-LacZ was not above background. This suggests that uMetE2 may be translated *in vivo* thereby potentially affecting riboswitch function. To investigate whether translation of uMetE2 had implications for MetE expression and B_12_-sensing, we mutated the start codon of uMetE2 to a non-start codon (UUG→UCG) in the context of the full-length MetE’-LacZ fusion (**Fig. 4B**; U124C). The construct was transformed into wild type (B_12_-producing) and Δ*cobK* (B_12_-deficient) *M. smegmatis* backgrounds to assess if B_12_-sensing was intact in the mutant. To our surprise, the mutation led to a >10-fold increase in MetE’-LacZ expression in both wild type and Δ*cobK* backgrounds, while the B_12_-dependent change was maintained (**Fig. 4C**).

This finding suggested that although the mutation dramatically altered overall expression, it did not impair B_12_ sensing. Conversely, deletion of the entire *umetE2* (segment spanning +115 to +188) led to complete loss of B_12_-sensing, while also increasing expression of MetE’-LacZ (**Fig. 4C**). In summary, these results suggest that translation of the aptamer-encoded uMetE2 contributes to the overall control of MetE expression. At this stage, we are unable to clarify whether it involves the uMetE2 peptide.

### ppe2 is preceeded by a riboswitch-controlled, translated leader

The results presented so far suggested multiple functional differences between the *metE* and *ppe2* switches. Therefore, to investigate in more detail the structure-function relationship of the *ppe2* switch, we performed inline probing of the *ppe2* leader (from +1 to +191) using a range of AdoB_12_ concentrations from 1 nM to 4 mM (**Fig. 5A**). The results indicated that cleavage of the *ppe2* riboswitch was modulated by AdoB_12_ in a dose-dependent manner, with strongly protected regions at G24-U29, G61-G64 and A108-G128 and hypercleavage at A131-C134 and U164/G165 (**Fig. 5A**). This result suggested that the sequences required for ligand binding, *i.e.,* the aptamer, were contained within the probed fragment. However, the affinity towards AdoB_12_ was considerably lower than that of the *metE* aptamer (K_d_ >400 μM (range = 426 μM – 767 μM) versus <60 μM) (**Supplementary Fig. 3**). We predicted the secondary structure of the ligand-bound *ppe2* riboswitch by using probing-derived folding constraints and a previously proposed KL interaction between L5 and L13 for this switch (31) (**Fig. 5C**). Similar to that of the *metE* riboswitch, the B_12_ binding pocket of the *ppe2* riboswitch was also enclosed in a four-way junction formed by the paired segments P3-P6. However, unlike the *metE* switch, the *ppe2* riboswitch contained a fused P1-P3 arm and a truncated P6 extension (**Fig. 5C**). The predicted structure suggests that the AdoB_12_-induced masking of positions 108-128 are likely due to a combination of base pairing and ligand contacts (**Fig. 6A** **& B**). Moreover, this structure replicates that of the *metE* switch where strong cleavage signals are observed immediately downstream of the B_12_ box (**Fig. 5A**; positions 131-134).

**Figure 5.**
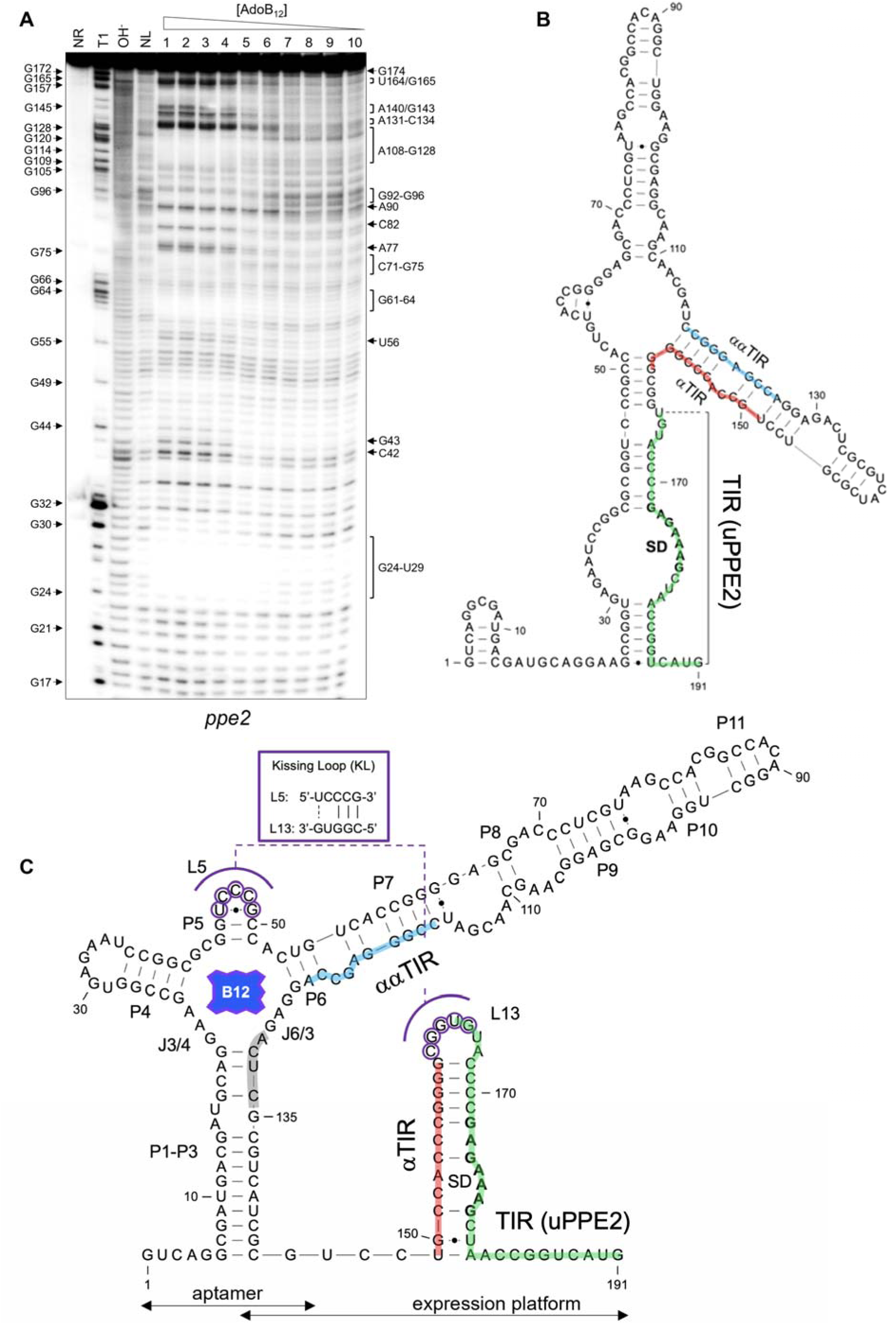
Structure of *ppe2* riboswitch. **A.** The cleavage pattern of the *ppe2* switch over a concentration gradient of AdoB_12_. Strongly modulated positions are indicated by labels on the right side of the gel. G positions are shown on the left. NR – no-reaction control; T1 – RNAse T1 ladder; OH^-^ – alkaline digest ladder; NL – no-ligand control. **B.** Predicted secondary structure of the Apo-form of the *ppe2* switch. The translation initiation region (TIR) including the SD sequence is highlighted in green, while the αTIR and ααTIR sequences are highlighted in red and blue, respectively. **C.** AdoB_12_-bound *ppe2* switch. Paired regions in the aptamer are labelled sequentially from P1 to P11. The “B_12_ box” located G121-G135 matches the consensus (68). The region of hypercleavage at A131-C134 is highlighted in grey. Bases of L5 that could potentially form a kissing loop (KL) with L13 are highlighted in purple and the base pairs between them are depicted in the inset.

**Figure 6.**
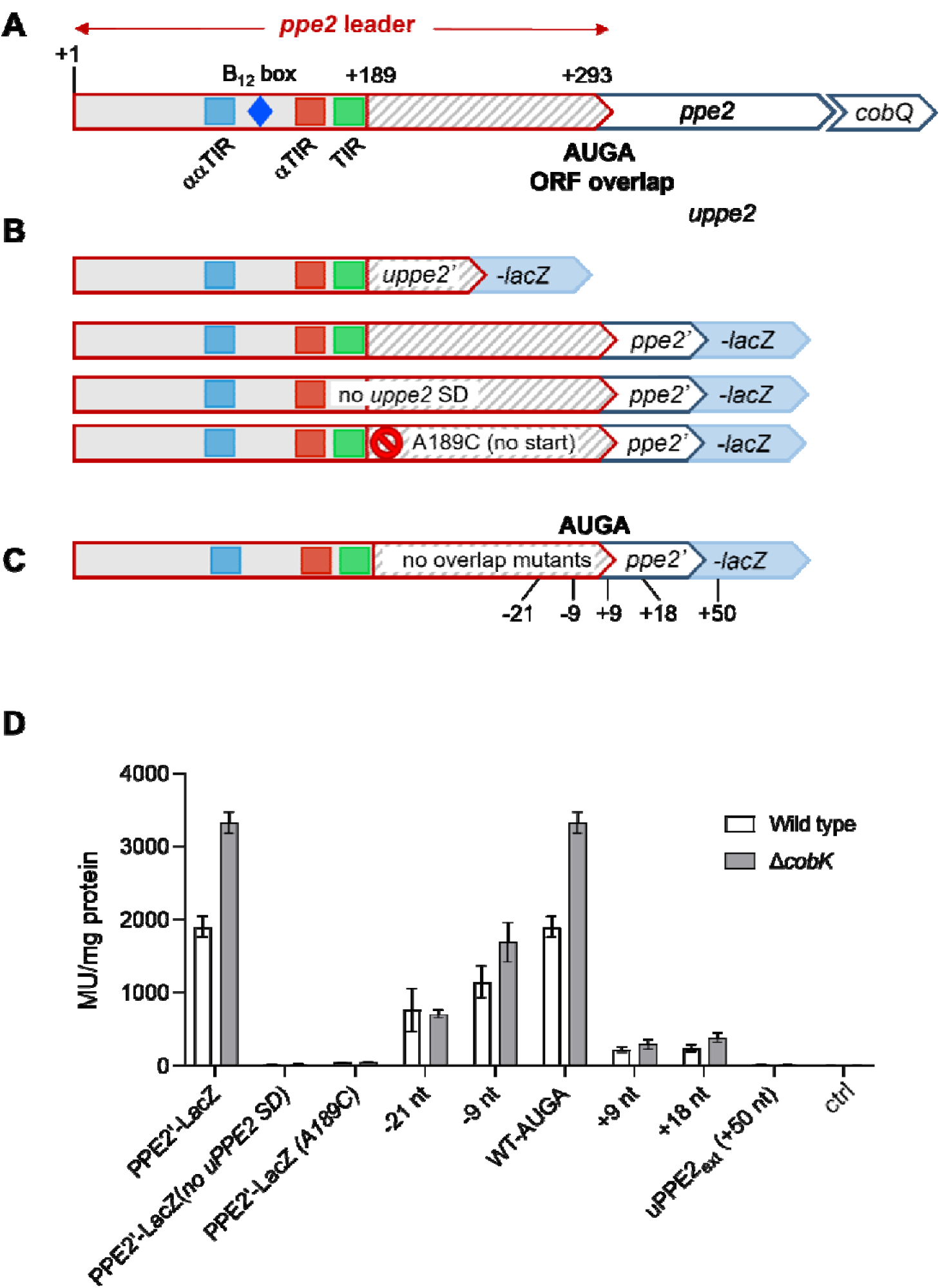
PPE2 are uPPE2 are co-regulated and co-translated. **A.** Outline of the *ppe2* operon indicating the regulatory elements, uPPE2 and shared AUGA overlap. **B.** Outline of *uppe2’-lacZ* and *ppe2’-lacZ* reporter fusions including non-translated uPPE2 variants (no SD and no start (A189C)). **C.** Locations of +50 extended uPPE2 stop codon and relocated stop codons for uPPE2. D. Result of β-gal reporter assays showing uPPE2-dependent translation of PPE2 in wild type and Δ*cobK* backgrounds (Miller units (MU)/mg total protein). Data represent mean ± standard deviation of three biological replicates. ctrl – no expression control.

The region immediately upstream of the annotated *ppe2* ORF is devoid of any obvious SD, but a potentially translated (21) and partially conserved uORF located between +189 and *ppe2* does appear to have a SD (**Fig. 5** **& Supplementary Fig. 6)**. In the predicted ligand-bound state, the uPPE2 TIR including this SD is partially masked by an αTIR stem, and together these elements form a hairpin that harbours a likely L5 pairing partner (L13, **Fig. 5C**).

In the Apo-structure prediction, the αTIR is sequestered by a purine-rich ααTIR sequence, while the SD and its downstream region are rendered more accessible for ribosome binding (**Fig. 5B**). Notably, the ααTIR preceeds and partially overlaps the B_12_ box (**Fig. 5C**). In summary, this structures suggests that the probed fragment contains both aptamer and expression platform, and we conclude that the *ppe2* switch presents as a very compact, translational “OFF” switch, which potentially controls the expression of uPPE2 in addition to *ppe2* and *cobQ*.

To gauge whether uPPE2 was in fact the first regulatory target in the *ppe2* operon, we made separate in-frame LacZ fusions of PPE2 and uPPE2 including the upstream riboswitch and 3 codons of uPPE2 for uPPE2’-LacZ and 13 codons of PPE2 for PPE2’-LacZ, to include the PPE motif located at residues 10-12 of PPE2; (**Fig. 6**). The constructs were transformed into *M. smegmatis* wild type and Δ*cobK* and the resulting β-gal assay indicated that uPPE2’-LacZ expression was higher than that of PPE2’-LacZ (**Table 1**). The partial conservation together with the high level of expression suggested that uPPE2 was indeed functional and the first target in the *ppe2* operon.

**Table 1:** β-gal activities of PPE2 and uPPE2 reporter strains

| Strain | PPE2'-LacZ | uPPE2'-LacZ | ctrl |
| --- | --- | --- | --- |
| Wild type | 1899.9 ± 143.0 | 2815.3 ± 120.4 | 3.9 ± 2.6 |
| $\Delta cobK$ | 3325.0 ± 147.9 | 4260.7 ± 208.2 | 5.4 ± 2.3 |

Data represent mean and standard deviation of three biological replicates

### Expression of PPE2 occurs via a uPPE2-PPE2 fusion protein

Hundreds of adjacent ORFs in *M. tuberculosis* have been found to share a 4-nt **N*UG****A* overlap between stop and start codons, which may affect translation of the downstream ORF (21, 65). As mentioned above, the annotated PPE2 ORF lacks a SD sequence, but it shares an **A*UG****A* stop/start overlap with the uPPE2 ORF (**Fig. 6A**). To ascertain whether translation of PPE2 relied on uPPE2, we introduced mutations to prevent translation initiation of uPPE2 in the fully-leadered PPE2’-LacZ fusion. In one construct, we eliminated the uPPE2 SD by replacing the purines with their complementary pyrimidines (*uPPE2 no SD*); in the other, the we changed the uPPE2 AUG start codon to the non-start CUG (uPPE2_A189C_) (**Fig 6B**). In both cases, we found that suppression of uPPE2 translation abolished PPE2 expression (**Fig. 6D**), suggesting that expression of PPE2 is strictly dependent on uORF translation. This may be due to Rho-dependent termination of transcription, as *RUT* sites become exposed in the absence of uPPE2 translation. Alternatively, it could be the result of direct translational coupling between the uPPE2 and PPE2 ORFs.

To determine whether uPPE2-dependent translation of PPE2 was due to Rho-dependent termination of transcription, we eliminated the native uPPE2 stop codon (A296U), while the PPE2 start codon remained intact. Thus, translation of uPPE2 continues until the next natural stop codon (50 nucleotides downstream, uPPE2_ext_), preventing Rho binding and termination. β-gal assays indicated that PPE2’-LacZ expression was also abolished in this context, suggesting that Rho-dependent termination did not account for the loss of expression (uPPE2_ext_ (+50 nt) **Fig. 6C** **&** **Fig. 6D**). To further dissect the mechanism underlying this phenomenon, which we suspected was directly linked to the stop-start overlap, we introduced stop codons in the **AUG**U/uPPE2_ext_ background at varying distances upstream (early stop) or downstream (late stop) of the PPE2 start codon (**Fig. 6C**). All constructs were expressed in *M. smegmatis* (wild type and Δ*cobK*) and β-gal activity determined. The results indicated that early stops led to a gradual reduction in PPE2’-LacZ expression and B_12_-sensing, which correlated with the stop-start separation distance (**Fig. 6D**). Conversely, shifting the stop codons downstream of the PPE2 start codon led to a more abrupt reduction in PPE2’-LacZ expression and B_12_ sensing (**Fig. 6D**). These results suggest that a tight stop/start overlap and a forward movement of the ribosome are necessary for efficient expression of the downstream ORF.

Translational coupling between overlapping ORFs can proceed via a Termination-Reinitiation (TeRe) mechanism leading to production of two separate polypeptides (21, 65), or via a frameshift without peptide release, which generates a fusion protein encoded by the two ORFs (72, 73). To determine which of these scenarios applied, we added an N-terminal FLAG-tag to uPPE2 in the PPE2’-LacZ reporter fusion (**Fig. 7A**). If translation coupling produced two distinct proteins, a signal corresponding to the expected size of FLAG-uPPE2 (6.3 kD) would be detected by western blotting; conversely, if the coupling resulted in a fusion protein, this would result in a signal corresponding to the size of the FLAG-uPPE2-PPE2’-LacZ fusion protein (123 kD). The constructs were expressed in *M. smegmatis* (wild type and Δ*cobK*) and cell extracts were prepared in parallel with cell extracts from the isogenic but non-FLAG-tagged reporter and from the background (no plasmid) strains for comparison. Western blotting using anti-FLAG antibodies indicated two strong signals around 130 kD, roughly corresponding to the expected size of the FLAG-uPPE2-PPE2’-LacZ fusion protein in the strains expressing the tagged uPPE2 (**Fig. 7B**; lanes 2 and 3), but not in the controls.

**Figure. 7.**
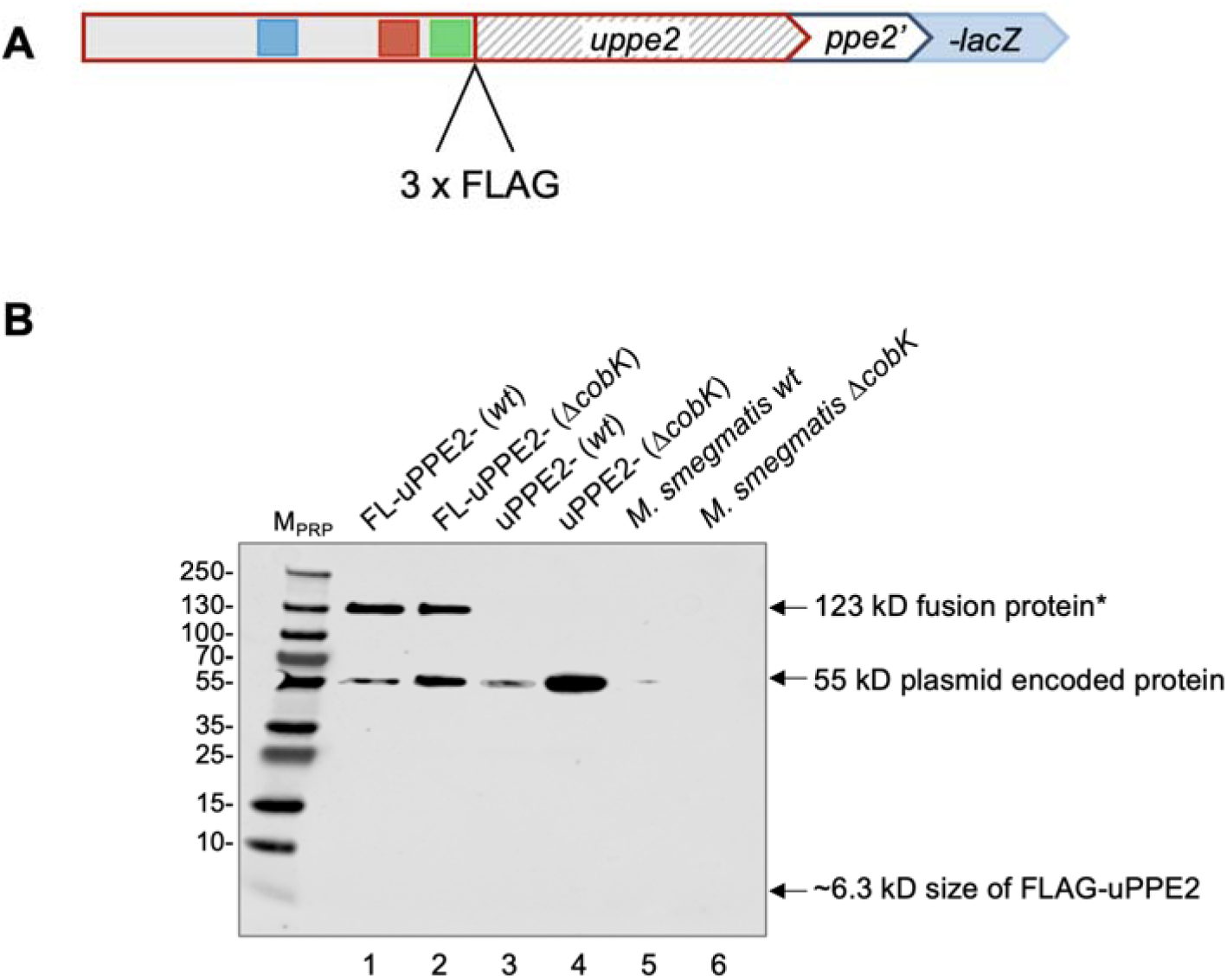
Expression of a uPPE2-PPE2 fusion protein. **A.** Schematic outlining the FLAG-tagged uPPE2 within the PPE2’-LacZ construct. **B.** Western blot detection of proteins from *M. smegmatis wildtype* (*wt*) or Δ*cobK* expressing FLAG-tagged uPPE2 (lanes 1 & 2), untagged uPPE2 (lanes 3 & 4) or without plasmid (lanes 5 & 6). Asterisk indicates likely 123-kD FLAG-tagged uPPE2-PPE2’-LacZ fusion protein. M_PRP_: Page Ruler Plus, protein ladder.

We observed an additional signal around 55 kD in all plasmid-carrying strains, but not in those without, suggesting that this was likely due to antibody cross-reactivity with an unspecific plasmid-encoded protein (**Fig. 7B**; lanes 4-7). More importantly however, there was no signal corresponding to the expected size of FLAG-uPPE2, suggesting that this peptide was either not made or it was rapidly degraded. In summary, these results suggest that translation of uPPE2 followed by a ribosomal frameshift are necessary for expression of PPE2, resulting in a uPPE2-PPE2 fusion protein.

## Discussion

In the current study, we have combined inline probing, structure prediction, biochemical and genetic approaches to compare the gene expression control mechanisms by two riboswitches in *M. tuberculosis*. Both switches are B_12_-sensing “OFF” switches, in agreement with previous observations for the *metE* switch and B_12_ riboswitches in general (26, 31, 34, 48, 49, 55, 57, 58, 74). Both riboswitches operate via B_12_-dependent masking and unmasking of the SD sequence, and in the case of *metE* this is accompanied by a massive reduction in mRNA levels, which is likely a result of Rho-dependent termination potentially in combination with mRNA degradation (21). The downregulation of *ppe2* mRNA was less pronounced. However, it is worth noting that the *ppe2* Apo-form starts with a stem-loop, which may increase mRNA stability compared to the ligand-bound form, which has several unpaired 5’ nucleotides (75).

Both riboswitches displayed features typical of B_12_ riboswitches such as the B_12_ box and kissing loop (KL), but there were also some unique features in both switches. For example, while the L5 half of the KL was identical in the two riboswitches, the interacting L13 differed slightly, although still providing similar base pairing. Moreover, the *metE* L13 overlapped with the proposed ααTIR, while *ppe2* L13 was flanked by the αTIR and TIR (**Fig. 3** **&** **Fig. 5**). Finally, the *metE* aptamer domain and the expression platform presented as two distinct domains, where the ααTIR was located well downstream of the B_12_ box and separated from the αTIR by a further ∼60 nucleotides (**Fig. 3**). This contrasted with the *ppe2* switch, where the overall size and distance between individual elements were much smaller. The ααTIR of the *ppe2* switch was located earlier in the switch in the P6/P7 extension, *i.e.*, upstream of the B_12_ box, indicating a substantial overlap between the aptamer domain and the expression platform. Moreover, the αTIR was located only ∼20 nucleotides further downstream of the ααTIR (**Fig. 5**).

In addition to the proposed structures, we demonstrate the presence of atypical and distinct regulatory uORF-related features that contribute to downstream gene expression in the two operons. uMetE2 is encoded within the P9-P10 extension of the aptamer upstream of the conserved B_12_ box. Mutating the uMetE2 start codon to eliminate translation significantly increased MetE expression, suggesting that uMetE2 is translated *in situ*, leading to suppression of MetE expression. Presumably, translation of uMetE2 and ligand-induced folding are mutually exclusive, but our results suggested that B_12_-sensing remains. Whether the suppression is a *cis* or *trans* effect, resulting from the peptide, remains unknown, but the fact that uMetE2 shows limited conservation (**Supplementary Fig. 4**) suggests that its function may be related to specific lifestyles of some members of the *M. tuberculosis* complex.

Another and rather unexpected feature was the indispensable relay between the B_12_aptamer and the *ppe2-cobQ(U)* operon via uPPE2. PPE2 is not expressed without translation of uPPE2, but requires suppression of the uPPE2 stop codon followed by a frameshift and the synthesis of a uPPE2-PPE2 fusion protein. To our knowledge, this is the first demonstration of stop codon suppression and translational coupling in *M. tuberculosis*. We are currently investigating how the sequence context surrounding the AUGA stop-start overlap might influence this mechanism.

We argue that the *ppe2* switch qualifies as a new sub-class of B_12_ riboswitches (Class IIc) based on the following considerations. Firstly, the *ppe2* switch does not belong to Class I since it has no P8-P12 extension. Secondly, even though it has the GGAA junctional motif at J3/4 similar to Class IIb switches, it lacks a corresponding UCU motif in the J6/3 junction, opposite (58). Moreover, the *ppe2* switch is neither unable to bind AdoB_12_, similar to Class IIa switches, nor AdoB_12_ selective, similar to Class IIb switches. Interestingly, positions C48-G50, which form the L5 of the *ppe2* switch, were protected from cleavage only in the presence of AdoB_12_ (**Supplementary Fig. 2**), implying that in this switch, the KL might be stabilised only by AdoB_12_ binding and not by MeB_12_, HyB_12_ or CNB_12_. Further structural analysis, such as crystalisation is required to obtain the full and true images of these switches in their ligand-bound and -unbound constellations (54, 55, 57).

It has been suggested that PE/PPE genes inserted and expanded at different genomic *loci* (52). Therefore, one can speculate that *ppe2* ‘invaded’ the locus of the cobalamin biosynthetic genes *cobQ/U,* which were initially under the regulatory control of an ancestral B_12_-riboswitch thereby giving rise to this extra and unconventional regulatory mechanism. This notion is supported to some extent by similar scenarios in the Mbox-*pe20*-*mgtC* locus and the recently identified PE-containing uORF in the *glyA2* locus (21).

The existence of B_12_ riboswitch classes with varying ligand selectivities within the same cell raises some interesting questions. It is assumed that *M. tuberculosis* relies on host-derived B_12_, but whether this is obtained by the pathogen as AdoB_12_ or MeB_12_ remains unknown. Moreover, nothing is known about potential pathways involved in converting the scavenged B_12_ isoform or precursor to the relevant riboswitch ligand or cofactor type. Therefore, understanding when and how *M. tuberculosis* utilises its multi-layered riboswitch complexity to sense a range of host niches will not only shed light on how habitats shape genomes, but also provide a deeper understanding of pathogen adaptation during the course of infection.

## Materials and Methods

### Strains and culture conditions

The *M. smegmatis* Δ*cobK* mutant was a gift from Professor Digby Warner. *E. coli* cells were grown either in Lysogeny Broth (LB) or on LB agar, with 250 μg/mL hygromycin, where appropriate. *M. smegmatis* cultures were grown either on Middlebrook 7H11 agar or Middlebrook 7H9 broth (Sigma-Aldrich) supplemented with glucose-salt solution (0.085% NaCl and 0.2% glucose). The *M. tuberculosis* H37Rv strain was cultured either in 7H9 broth supplemented with 10% ADC (Remel) or on 7H11 agar supplemented with 10% Middlebrook OADC (Becton-Dickinson). Cloning was performed in *E. coli* DH5α electrocompetent cells (New England Biolabs). Plasmids were extracted with Qiagen miniprep kits (Qiagen GmbH) and verified via Sanger sequencing (Source Biosciences). Transformation in mycobacteria was performed by electroporation using the exponential protocol settings (2.5 kV; 25 μF; 1000Ω). For antibiotics selection and blue-white screening of mycobacteria, 7H11 plates contained 50 μg/mL hygromycin and 50 μM 5-bromo-4-chloro-3-indolyl-β-D-galactopyranoside (X-gal) (Thermo Scientific). Unless specified, *M. tuberculosis* H37Rv culture was supplemented with 10 μM exogenous adenosylcobalamin (Thermo Scientific). Methylcobalamin, and hydroxocobalamin were purchased from Cambridge Bioscience whereas cyanocobalamin was obtained from Sigma-Aldrich.

### Oligonucleotides, plasmids, and cloning

Oligos and plasmids used in this study are listed in **Supplementary Table 1**. DNA oligos longer than 100 bp were purchased as geneBlocks fragments from Integrated DNA Technologies. Other oligos and primers were purchased from Thermofisher Scientific. Reporter constructs were generated by inserting target sequences using either Gibson assembly or restriction cloning between the *Hin*dIII and *Nco*I sites of pIRaTE2020 (21), to produce in-frame translational fusions with LacZ. The 5’ edges of *umetE’-lacZ* and *umetE4’-lacZ* fusions were +87 and 159, respectively, relative to the TSS (+1) of the *metE* leader; the 5’ boundaries of other *lacZ* fusions are specified in the relevant sections of the manuscript. Nucleotide substitution or deletion mutations were designed on the NEBaseChanger online tool and TOPO-cloned using the Q5 site-directed mutagenesis kit (New England Biolabs) according to the standard protocol.

### RNA extraction and northern blot analysis

*M. tuberculosis* H37Rv cultures were rapidly chilled by directly mixing with ice and pelleted by centrifugation. RNA was isolated using the RNAPro Blue kit (MP Biomedicals) according to the manufacturer’s protocol. RNA concentration and quality were evaluated on a Nanodrop 2000 spectrophotometer (Thermo Scientific). For northern blot analysis, 10 μg total RNA was separated on denaturing 8% polyacrylamide gel and transferred on a blotting paper for detection, as previously described (75). RNA probes were synthesised using the mirVana miRNA probe synthesis kit (Ambion) and radiolabelled using 133 nM ^32^P α-UTP (3000 Ci/mmol; Hartmann Analytic GmbH), with unlabelled UTP added to achieve a final concentration of 3 μM. RNA fragment signals were developed on radiosensitive screens and visualised on a Typhoon FLA 9500 phosphorimager (GE Healthcare).

### Quantitative real-time PCR

The RNA used for quantitative real-time PCR (qRT-PCR) was isolated from cultures supplemented with 10 μg/mL exogenous AdoB_12_, as reported by others (40, 48). qRT-PCR was performed on the QuantStudio 6 real-time PCR system (Applied Biosystems) using cDNA synthesised from 0.5 μg DNAse-treated RNA using the Superscript IV Reverse Transcriptase kit (Invitrogen) and random hexamers. Each 20-μL PCR reaction contained 1× Fast SYBR Green Master Mix (Applied Biosystems), 200 nM forward and reverse primers and 5 μL of 100× diluted cDNA or *M. tuberculosis* genomic DNA standards. The leader amplicons of *metE* covered the nucleotides from -93 to +22 relative to the *metE* coding sequence, while the *ppe2* leader amplicon covered from -121 to -2 relative to the *ppe2* coding sequence. The primers for the *metE* coding amplicon amplified from +355 to +460 relative to the start of the *metE* coding sequence, while the *ppe2* coding amplicon covered the region from +715 to +825 relative to the start of the *ppe2* coding sequence. The *ppe2-cobQ* junction amplicon covered the segment from +1569 to +1826 relative to the start of the *ppe2* coding sequence. Relative gene expression was determined as a ratio of the level of target mRNA to that of 16S rRNA. All data were graphed and analysed using GraphPad Prism software for Mac OS, version 9.0 (www.graphpad.com).

### Beta-galactosidase (**β−**gal) activity assay

Protein expression levels were assessed using the beta-galactosidase (β−gal) activity assay as previously described (75). Briefly, 10-mL cell cultures were centrifuged, and the cell pellet washed thrice in Z-buffer (60 mM Na_2_HPO_4_, 40 mM NaH_2_PO_4_, 10 mM KCl, 1mM MgSO_4_) prior to lysing in a FastPrep bio-pulveriser (MP Biomedicals). The protein concentration in the cell lysate was determined using the BCA kit (Thermo Scientific) according to the manufacturer’s instructions. The level of *lacZ* expression was calculated in Miller units (M.U.) per milligram of protein. All data were graphed and analysed using GraphPad Prism software, version 9.0 (www.graphpad.com).

### FLAG-tagging and western blot analysis

A triple FLAG-tag was inserted to the N-terminus of uPPE2 in the *ppe2-lacZ* reporter construct (pTKSW-*ppe2*; **Supplementary Table 1**) using Q5 site-directed mutagenesis (New England Biolabs) with the forward primer (5’-CATGATATCGACTACAAAGACGATGACGACAAGACGCTCCAAACCTTGTCT-3’) and reverse primer (5’-ATCCTTATAATCACCATCATGATCCTTATAATCCATGACCGGTTAGCTTTC-3’). The construct was transformed into *M. smegmatis* wild type and Δ*cobK* strains. Then, 100 mL of log-phase (OD_600_ ∼ 0.6) cultures were pelleted, resuspended in Z buffer, and lysed in FastPrep machine as described in the above protocol for the beta-galactosidase (β−gal) activity assay. For western blotting, 20 μL of the supernatant was loaded onto a 4-20% Mini-PROTEAN Tris-glycine gradient gel (Bio-Rad) and proteins separated by SDS-PAGE were transferred to a PVDF membrane. The FLAG-tagged protein was detected by incubating the milk powder-blocked membrane with mouse monoclonal anti-FLAG primary antibody (Sigma-Aldrich) diluted at 1:1000, followed by incubation with peroxidase-conjugated polyclonal goat anti-mouse IgG (Jackson ImmunoResearch) diluted at 1:1000. Signals were visualised on a Li-Cor Odyssey Fc Imager (Licor).

### In-line probing

In-line probing was performed as described by Regulski and Breaker (66). Riboswitch RNA was transcribed using the Megascript T7 High Yield Transcription Kit (Invitrogen) from amplicons generated by PCR using the primers listed in **Supplementary Table 1**. Transcribed RNA was extracted from denaturing 8% polyacrylamide gel in 500 μL crush-and-soak buffer (0.5 mM sodium acetate pH 5.2, 0.1% SDS, 1 mM EDTA pH 8.0, 30 μL acid phenol:chloroform) and precipitated in ethanol. The yield and purity of the transcribed product were analysed on a Nanodrop 2000. Dephosphorylation was done using calf intestinal alkaline phosphatase (1 U/μL) (Life Technologies). Each 25-μL 5’ end-labelling reaction contained 10 pmol RNA, 133 nM ^32^P γ-ATP (6000Ci/mmol; Hartmann Analytic GmbH), 1 μM unlabelled ATP, and 25U T4 polynucleotide kinase (10 U/μL) (New England Biolabs). The radiolabelled RNA was purified by PAGE and resuspended in 40 μL nuclease-free water. For in-line probing, reactions contained 2 μL radiolabelled RNA (∼25 nM), 1× in-line reaction buffer (50 mM Tris-HCl pH 8.3, 20 mM MgCl_2_, 100 mM KCl), and the desired concentration of AdoB_12_, MeB_12_, HyB_12_ or CNB_12_ in 20-μL total volume. Reactions were incubated on a heat block maintained at 30 °C for 20 hours and quenched with an equivalent volume of 2× colourless gel-loading solution (10 M urea, 1.5 mM EDTA pH 8.0). In-line reaction products were separated by denaturing 6-10% polyacrylamide gel electrophoresis (PAGE) at 45W. The gels were dried and exposed to radiosensitive screens and the data were collected on a Typhoon FLA 9500 (GE Healthcare). The dissociation constants (K_d_) of the riboswitches were calculated from the inline probing data in Fig. 3A and Fig. 5A, by plotting the fraction of RNA cleaved at ligand-sensitive (cleaved) sites against the logarithm of AdoB_12_ concentration, using the formula described in (66). In Graphpad prism (version 9.0), the data were fitted using a log(agonist) vs. response curve to obtain K_d_ values (*metE* riboswitch: K_d_ = 16.1 μM (95% confidence interval 5-57μM); *ppe2* riboswitch: K_d_ = 572.1μM (95% confidence interval 426-767μM).

### Sequence alignments

The DNA sequences of *M. tuberculosis metE* and *ppe2* leaders stretching from ∼40 nucleotides upstream of the TSS to the first codon of the downstream annotated ORF were used as the input query for nucleotide alignment on the NCBI BLAST tool (76). Matches of >95% identity in representative mycobacteria were extracted and their TSS located by examining their respective -10 elements. The 5’ ends of the shortlisted sequences were trimmed to only -5 nucleotides relative to the TSS. The DNA sequence was converted to RNA prior to alignment using t-coffee with default settings (77). Amino acid sequences of *uPPE2* were similarly aligned using t-coffee default settings for protein alignment (77). The resulting clustalw format alignment files were downloaded and edited using the desktop version of Jalview (version 2.11.2.5) (78).

### RNA secondary structure prediction and visualisation

Target RNA sequences and matching folding constraints were loaded on the RNAstructure web servers (67), and the structure prediction software ran using default RNAstructure tools (version 6.4). A MaxExpect file containing a CT-formatted structure was downloaded and converted to a dot-bracket-formatted file, which was used to render the 2D secondary structure using the web-based RNA2drawer app (79).

## Acknowledgments

The authors thank Digby Warner for supplying the *M. smegmatis* Δ*cobK* strain, Finn Werner for critically reading the manuscript, Fabian Blombach and Winnie Sambu for their helpful advice and insightful discussions.

## Author Contributions

All authors contributed substantively to the work presented in this paper. TK and KBA designed the study. TK, PP, DB, AD, ZP conducted experiments. TK, PP, KBA performed data analysis and wrote the manuscript. All authors discussed the findings and their implications, and commented on the manuscripts at various stages.

