## Supplementary figures for "*Mycobacterium tuberculosis* employs atypical and different classes of B_12_ switches to control separate operons"

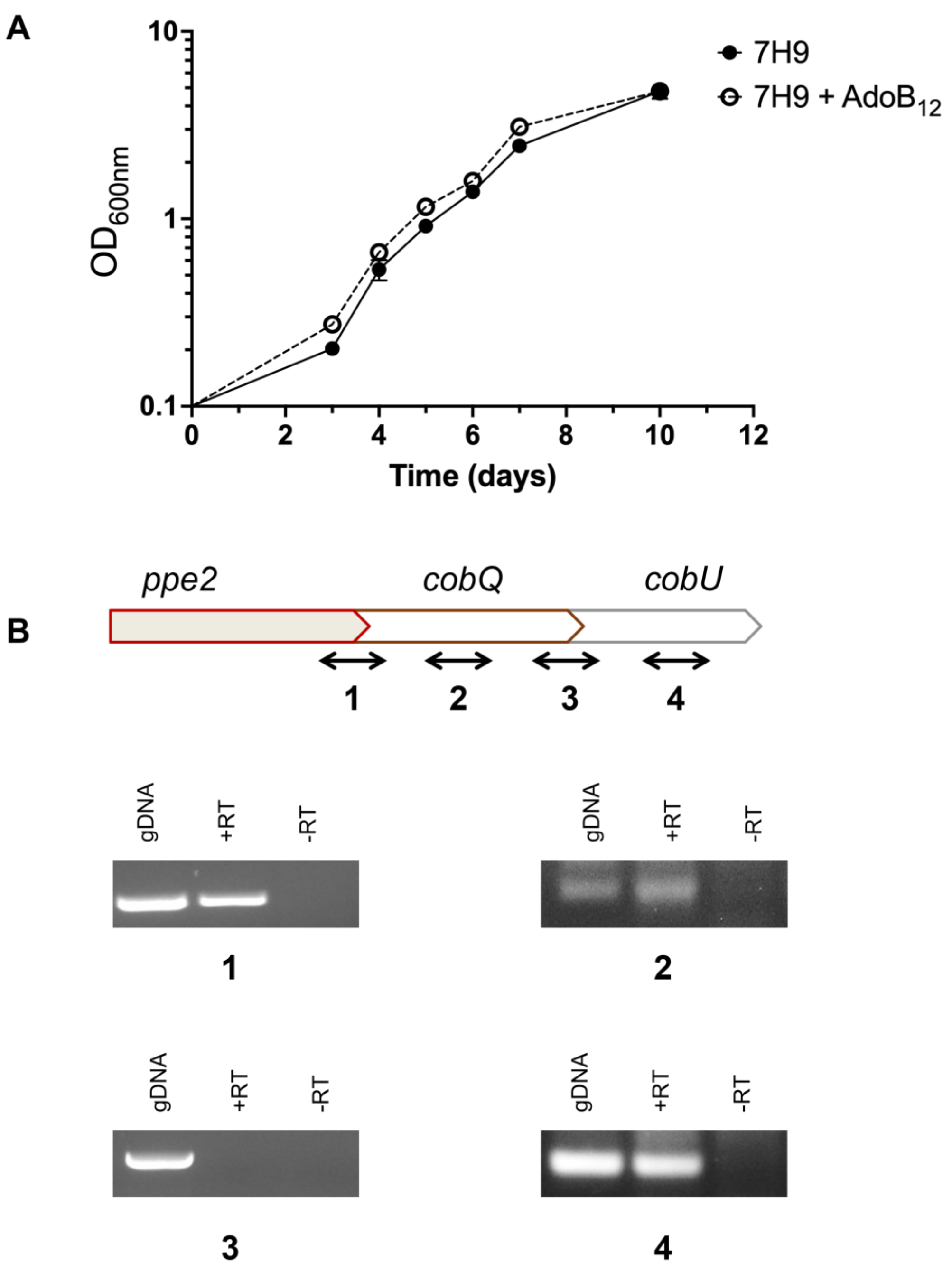


**Supplementary Figure 1**. **A**. Growth curve of *M. tuberculosis* H37Rv in 7H9/ADC media supplemented with $AdoB_{12}$ or without supplement. The data are plotted as mean ± standard deviation of triplicate cultures. **B**. A schematic showing the relative positions of RT-PCR primers (not to scale) targeting the *ppe2-cobQ* junction (1), *cobQ* coding region (2), *cobQ-cobU* junction (3), and *cobU* coding region (4). RT-PCR was performed using cDNA generated from RNA isolated in *M. tuberculosis* H37Rv cultures without supplement.

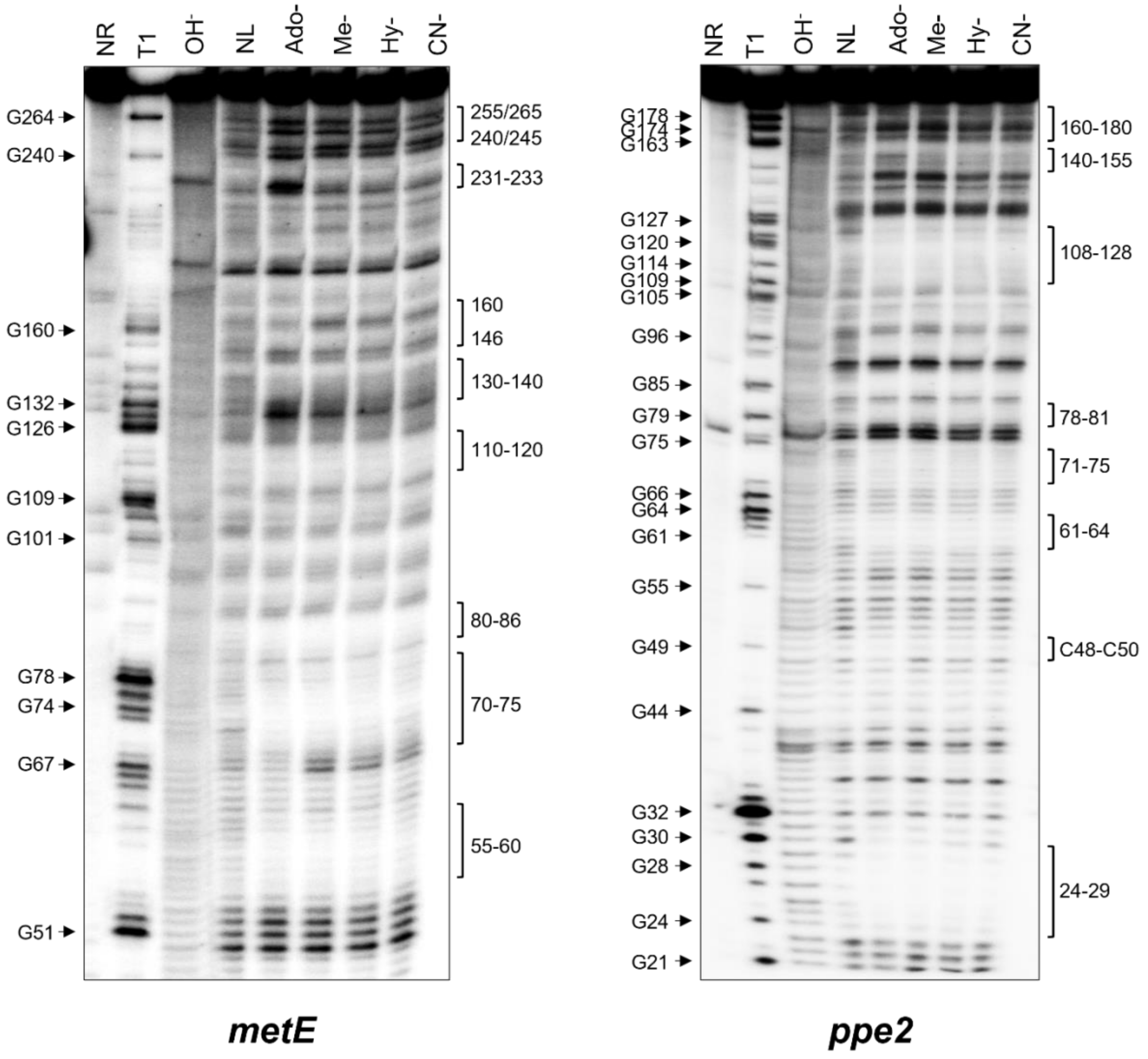


**Supplementary Figure 2**. Inline probing reveals differences in sensitivities of *metE* and *ppe2* riboswitches to common $B_{12}$ isoforms. Reactions contained 1 mM adenosylcobalamin (Ado-), Me- (methylcobalamin), Hy- (hydroxocobalamin), or CN- (cyanocobalamin). Regions showing prominent ligand-induced modulations are indicated with brackets to the right of each gel. G positions are based on RNAse T1 digestion. NR – no-reaction control; T1 – RNAse T1 ladder; $OH^-$ – alkaline hydrolysis ladder; NL – no-ligand.

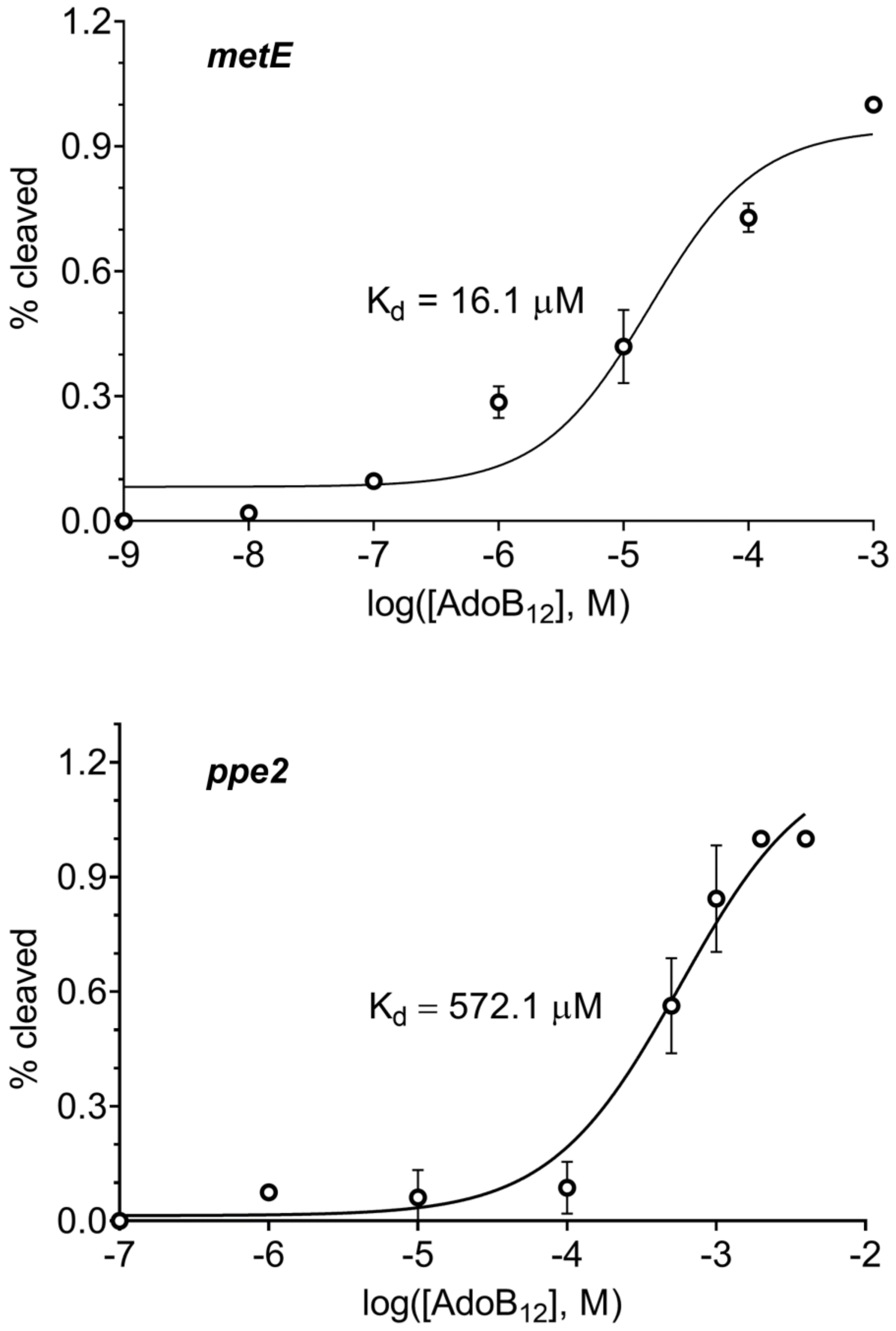


**Supplementary Figure 3**. Dose response curves (log(agonist) vs. response) for $AdoB_{12}$ binding by *metE* and *ppe2* riboswitches. The dissociation constants ($K_d$) of the riboswitches calculated from the inline probing data in Fig. 3A and Fig. 5A (in the main manuscript) and are indicated. The 95% confidence interval of the $K_d$ values of *metE* and *ppe2* riboswitches are 5.0-56.5 μM and 425.7-767.8 μM, respectively.

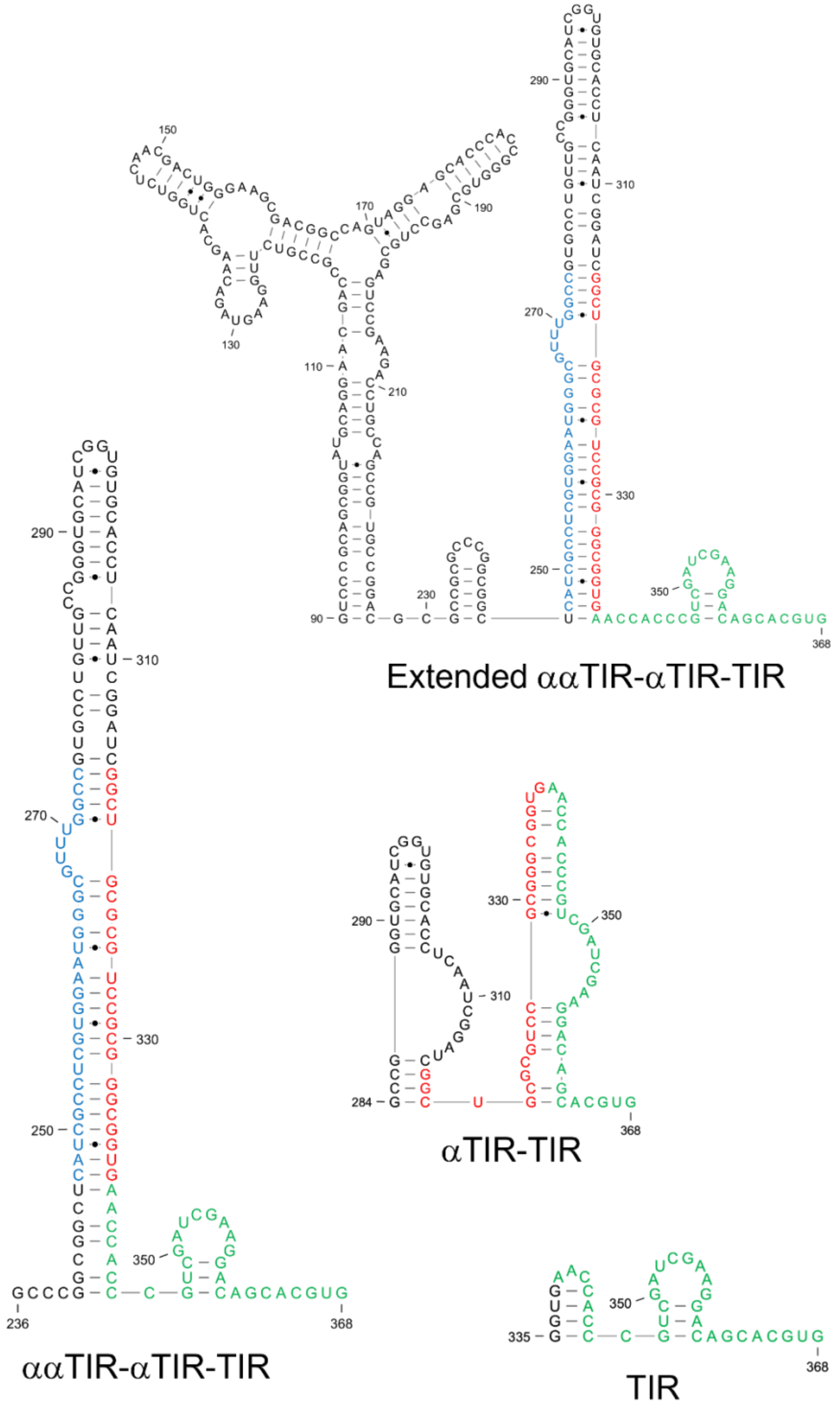


**Supplementary Figure 4**. Folding structures composed of highly probable base pairs for constructs of gradually increasing 5’ ends from the *metE* start codon showing translation control elements: TIR -translation initiation region; αTIR – sequestrator of TIR; ααTIR – anti-sequestrator. The first base in each structure is numbered relative to the *metE* TSS (+1).

### Supplementary Figure 5

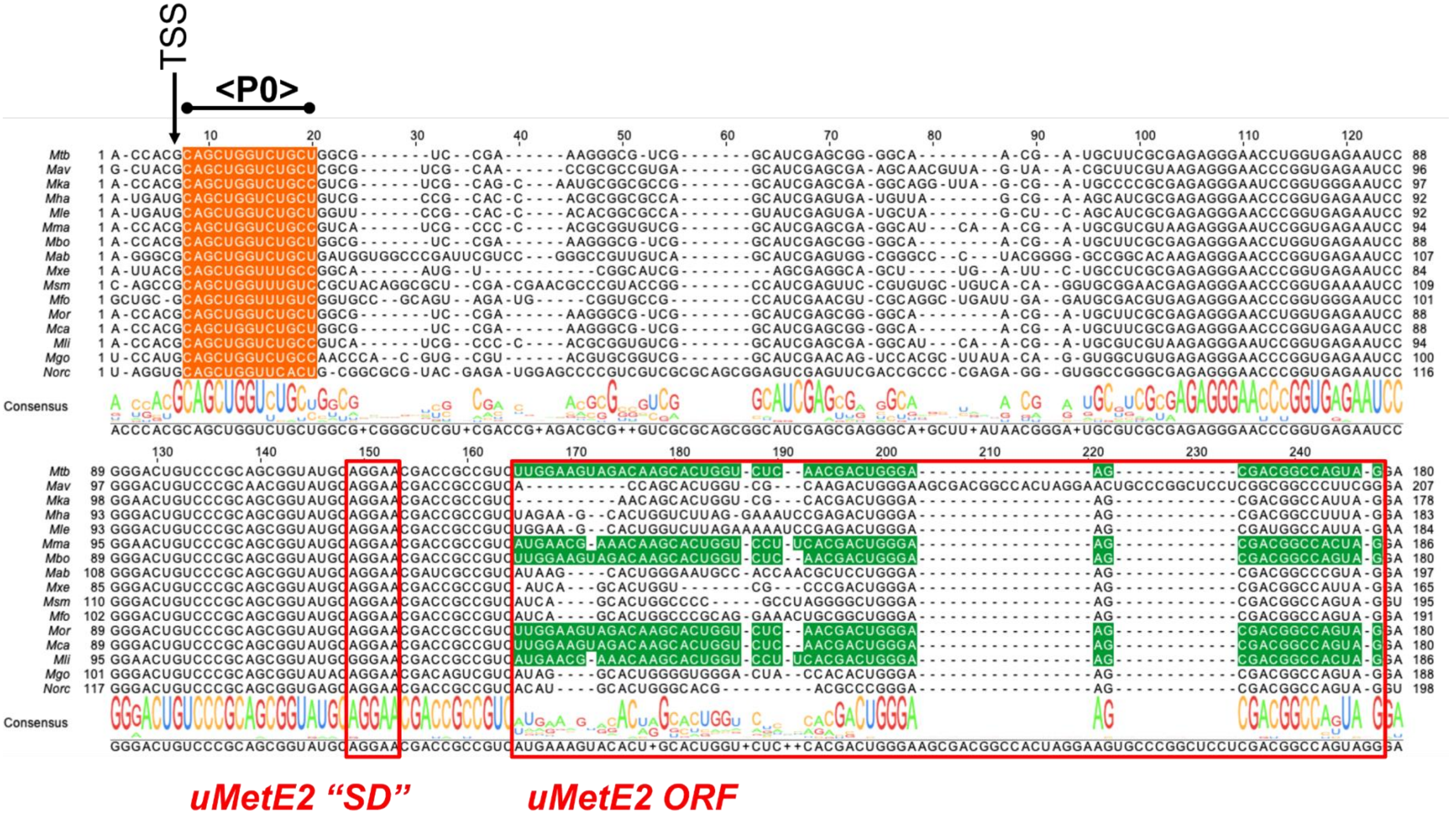

**Supplementary Figure 5**. Nucleic acid sequence alignment of the first 175 bases in the *metE* riboswitch aptamer. The ultra-conserved P0 element immediately downstream of the TSS is highlighted in orange. The SD-like sequence of uMetE2 is shown by a red box and the reading frame is highlighted in green and a red border enclosing the region occupied by the uORF. *Mtb – M. tuberculosis; Mav – M. avium; Mka – M. kansasii; Mha – M. haemophilum; Mle – M. leprae; Mma – M. marinum; Mbo – M. bovis; Mab – M. abscessus; Mxe – M. xenopi; Msm – M. smegmatis; Mfo – M. fortuitum; Mor – M. orygis; Mca – M. canettii; Mli – M. liflandii; Mgo – M. goodii; Norc – Nocardia sp.* strain CS682.

**Supplementary Figure 6**

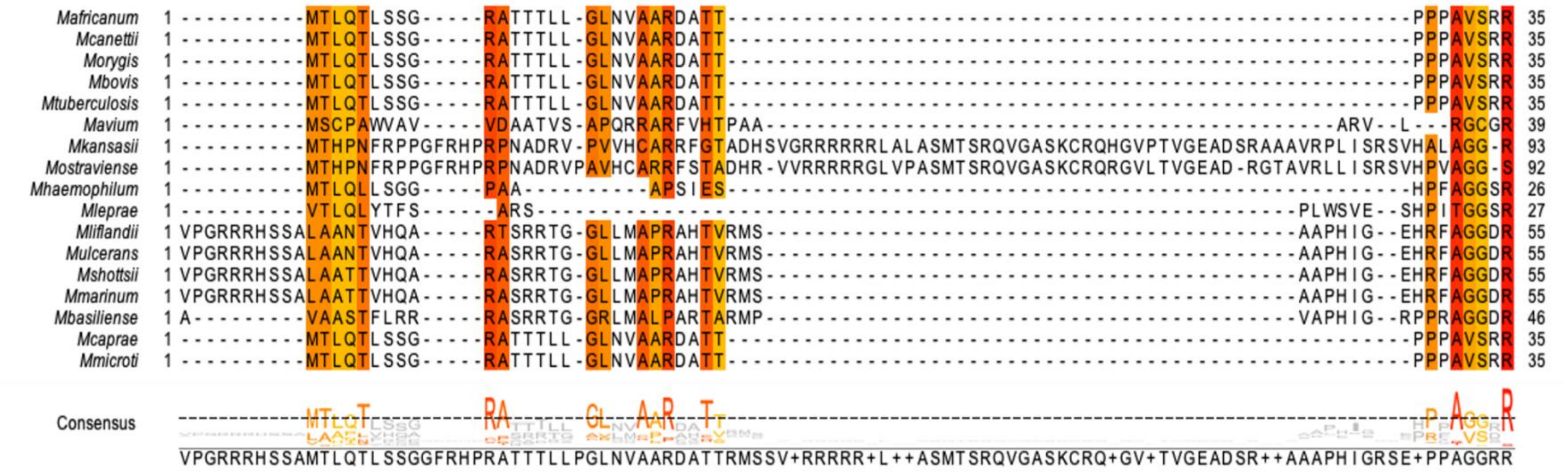

**Supplementary Figure 6**. Protein sequence alignment showing conserved uPPE2 residues in representative mycobacteria. Amino acids with > 50% identity are coloured, with red showing the highest conservation.
