## Supplementary table 1 for "*Mycobacterium tuberculosis* employs atypical and different classes of B_12_ switches to control separate operons"

### **Supplementary Table 1: Strains, plasmids, oligos, primers, and gene blocks fragments**

| **Strains** | | |
| --- | --- | --- |
| **Strain** | **Notes** | **Reference(s)** |
| *E. coli* DH5α | Chemically competent *E. coli* cells capable of high efficiency transformation | New England Biolabs |
| *M. smegmatis* mc^2^155 | Wild type strain of transformation-efficient *M. smegmatis* | (1) |
| *M. smegmatis* Δ*cobK* | *M. smegmatis* mutant with unmarked deletion of *cobK* (MSMEG_3875) | (2) |
| *M. tuberculosis* H37Rv | Laboratory-adapted strain of *M. tuberculosis* | (3) |

| **Plasmids** | | |
| --- | --- | --- |
| **Plasmid name** | **Description** | **Reference(s)** |
| pIRaTE2020 | A derivative of pIRaTE containing *Hin*dIII and *Nco*I restriction cloning sites and a PCL1 core promoter driving the expression of a *lacZ* reporter; hygromycin selectable | (4, 5) |
| pTKSW-*ppe2* | A derivative of pIRaTE2020 containing the leader and the first 13 codons of *ppe2* fused in-frame to *lacZ* | This study |
| pTKSW-*metE* | A derivative of pIRaTE2020 containing the leader and the first 2 codons of *metE* fused in-frame to *lacZ* | This study |
| pTKSW-*uPPE2* | A derivative of pTKSW-*ppe2* containing a site-directed deletion of the region from +198 to +332 of the *ppe2* leader, which generates an in-frame fusion of the leader of *uPPE2* and its first three codons to *lacZ* | This study |
| pTKSW-*ppe2_A296U_* | A derivative of pTKSW-*ppe2* containing an *UGA→UGU* mutation in the stop codon of *uPPE2* | This study |
| pTKSW-*FLAG-uPPE2* | A derivative of pTKSW-*ppe2* containing a site-directed insertion of a triple FLAG-tag sequence (GATTATAAGGATCATGATGGTGATTATAAGGATCATGATATCGACTACAAAGACGATGACGACAAG) to the N-terminus of uPPE2 | This study |

| **Oligos and Primers** | | | |
| --- | --- | --- | --- |
| **Oligo ID** | **5’ → 3’ sequence** | **Description** | **Reference(s)** |
| 1.48 | GTCCCATTCCGAACCCGGAAGCTAAGCCTGCCAGCGCCTGTCTC | Northern blot riboprobe targeting the 5S rRNA transcript | (4) |
| 3.07 | AAGAAGCACCGGCCAACTAC | Forward primer for qRT-PCR targeting the 16S rRNA transcript | (4) |
| 3.08 | TCGCTCCTCAGCGTCAGTTA | Reverse primer for qRT-PCR targeting the 16S rRNA transcript | (4) |
| 10.73 | GGCGTCCGAAAGGGCGTCGGCATCGAGCGGGGCAACGATGCCTGTCTC | Northern blot riboprobe targeting the *metE* leader transcript | This study |
| 10.74 | GGAGCGACCCTCGTAAGCCACGGCCACAGGCTGGAAGGCGCCTGTCTC | Northern blot riboprobe targeting the *ppe2* leader transcript | This study |
| 11.12 | TAATACGACTCACTATAGGGCAGCTGGTCTGCTGGCGTC | Forward primer to PCR-amplify the template used to transcribe a 345-nt transcript for inline probing of the *metE* switch | This study |
| 11.11 | GGTGGTTTCACCGCCCGCGGA | Reverse primer to PCR-amplify the template used to transcribe a 345-nt transcript for inline probing of the *metE* switch | This study |
| 11.07 | TCGCGAGAGGCCCCCTGGTGAGAATCCG | Forward primer used to generate 67-69GAA>CCC mutation in the pTKSW-*metE* construct via SDM |  |
| 11.08 | AGCATCGTTGCCCCGCTC | Reverse primer used to generate 67-69GAA>CCC mutation in the pTKSW-metE construct via SDM |  |
| 11.09 | CCGGGACTGTAAAGCAGCGGTATG | Forward primer used to generate 92-94CCC>AAA mutation in the pTKSW-*metE* construct via SDM |  |
| 11.10 | ATTCTCACCAGGTTCCCTC | Reverse primer used to generate 92-94CCC>AAA mutation in the pTKSW-metE construct via SDM |  |
| 11.20 | TAATACGACTCACTATAGGGTCAGGCGATGACGATGCAG | Forward primer to PCR-amplify the template used to transcribe a 191-nt transcript for inline probing of the *ppe2* switch | This study |
| 12.26 | CATGACCGGTTAGCTTTCT | Reverse primer to PCR-amplify the template used to transcribe a 191-nt transcript for inline probing of the *ppe2* switch | This study |
| 11.48 | CTAACCGGTCCTGACGCTCCAAA | Forward primer used to generate *uPPE2_A189C_* in the pTKSW-*ppe2* construct via site-directed mutagenesis (SDM) | This study |
| 11.49 | CTTTCTCGGGGTACACCG | Reverse primer used to generate *uPPE2_A189C_* in the pTKSW-*ppe2* construct via SDM | This study |
| 11.80 | Phos/AGCTTACTGTCCCGCAGCGGTATGCAGGAACGACCGCCGTCTTGGAAGTAGACAAGCACTGGTCTCC | Used to generate the *uMETE2-lacZ* (SD only) construct by oligo annealing and restriction cloning | This study |
| 11.81 | Phos/CATGGGAGACCAGTGCTTGTCTACTTCCAAGACGGCGGTCGTTCCTGCATACCGCTGCGGGACAGTA | Used to generate the *uMETE2-lacZ* (SD only) construct by oligo annealing and restriction cloning | This study |
| 12.01 | GCCGCCGATGTCCGCCCCGATC | Forward primer used to generate pTKSW-*ppe2_A296U_* via SDM | This study |
| 12.02 | TGACCGCCGGCGGAGGTG | Reverse primer used to generate pTKSW-*ppe2_A296U_* via SDM | This study |
| 12.18 | CCATGGATGATCCCGTCG | Forward primer to generate pTKSW-*uPPE2* via SDM | This study |
| 12.19 | GAGCGTCATGACCGGTTA | Reverse primer to generate pTKSW-*uPPE2* via SDM | This study |
| 12.20 | Phos/AGCTTAGCGACGGCCAGTAGGAGCACCCACCGGGTGCGAGCCTGCGAGCC | Used to generate the *uMETE4-lacZ* (SD only) construct by oligo annealing and restriction cloning | This study |
| 12.21 | Phos/CATGGGCTCGCAGGCTCGCACCCGGTGGGTGCTCCTACTGGCCGTCGCTA | Used to generate the *uMETE4-lacZ* (SD only) construct by oligo annealing and restriction cloning | This study |
| 16.58 | ACCGCCGTCTCGGAAGTAGAC | Forward primer used to generate *metE_U124C_* in the pTKSW-*metE* construct via SDM | This study |
| 16.59 | CGTTCCTGCATACCGCTG | Reverse primer used to generate *metE_U124C_* the pTKSW-*metE* construct via SDM | This study |
| 12.43 | CCATGGATGATCCCGTCGTTTTAATTAG | Forward primer used to generate pTKSW-*uMETE2* via SDM | This study |
| 12.44 | ACTGGCCGTCGCTTCCCA | Reverse primer used to generate pTKSW-*uMETE2* via SDM | This study |
| 12.52 | GACGACACCTTGACCGGCGGTCAGCC | Forward primer used to introduce a stop codon at -21 nt relative to *ppe2* start codon in the pTKSW-*ppe2_A296U_* construct via SDM | This study |
| 12.53 | GCGTCGCGAGCAGCCACG | Reverse primer used to introduce a stop codon at -21 nt relative to *ppe2* start codon in the pTKSW-*ppe2_A296U_* construct via SDM | This study |
| 12.54 | TGAAGCCGCCGATGTCCGCCC | Forward primer used to introduce a stop codon at -9 nt relative to *ppe2* start codon in the pTKSW-*ppe2_A296U_* construct via SDM | This study |
| 12.55 | GACCGCCGGCGGAGGTGTC | Reverse primer used to introduce a stop codon at -9 nt relative to *ppe2* start codon in the pTKSW-*ppe2_A296U_* construct via SDM | This study |
| 12.56 | ATGTCCGCCCTGATCTGGATGGCTTCGC | Forward primer used to introduce a stop codon at +9 nt relative to *ppe2* start codon in the pTKSW-*ppe2_A296U_* construct via SDM | This study |
| 12.57 | CGGCGGCTGACCGCCGGC | Reverse primer used to introduce a stop codon at +9 nt relative to *ppe2* start codon in the pTKSW-*ppe2_A296U_* construct via SDM | This study |
| 12.58 | GATCTGGATGACTTCGCCCCCAGAGGTGC | Forward primer used to introduce a stop codon at +18 nt relative to *ppe2* start codon in the pTKSW-*ppe2_A296U_* construct via SDM | This study |
| 12.59 | GGGGCGGACATCGGCGGC | Reverse primer used to introduce a stop codon at +18 nt relative to *ppe2* start codon in the pTKSW-*ppe2_A296U_* construct via SDM | This study |
| 12.60 | GTGTACCCCGTCTTTCCTAACCGGTCATGACGCTCCAAACCTTGTC | Forward primer used to generate *uPPE2_flipped SD_* in the pTKSW-*ppe2* construct via SDM | This study |
| 12.61 | CGCCCCGGGTGGCAGGAC | Reverse primer used to generate *uPPE2_flipped SD_* in the pTKSW-*ppe2* construct via SDM | This study |
| 14.68 | GCCGGATTTGTATTAGACTAAGCTTGAGTAGGAGATTTTCACCTCCTTTCCTTCCTACCATGGATG ATCCCGTCGTTTTACA | Used to generate a no-expression control via oligo annealing and Gibson assembly of an insert containing purine-pyrimidine substitutions in the *lacZ* SD sequence of pIRaTE2020 | (5) |
| 14.69 | TGTAAAACGACGGGATCATCCATGGTAGGAAGGAAAGGAGGTGAAAATCTCCTACTCAAGCTTAGTCTAATACAAATCCGGC | Used to generate the no-expression control via oligo annealing and Gibson assembly of an insert containing purine-pyrimidine substitutions in the *lacZ* SD sequence of pIRaTE2020 | (5) |
| 16.03 | TCGCTGGGTAATCCGCTAAC | Forward primer for qRT-PCR targeting *ppe2* ORF (downstream amplicon) | This study |
| 16.04 | GAATGCGAAGGTTTGCGACA | Reverse primer for qRT-PCR targeting *ppe2* ORF (downstream amplicon) | This study |
| 16.01 | TGGTTCGACACCAACTACCA | Forward primer for qRT-PCR targeting *metE* ORF (downstream amplicon) | This study |
| 16.02 | GCCCTAACGCCTCTTTGAGT | Reverse primer for qRT-PCR targeting *metE* ORF (downstream amplicon) | This study |
| 16.05 | ATCGAGTTGTTGGACATGTTC | Reverse primer for PCR (and qRT-PCR) targeting the *ppe2-cobQ* junction | This study |
| 16.06 | CGCAGGACTGATCACGTTA | Forward primer for PCR (and qRT-PCR) targeting the *ppe2-cobQ* junction | This study |
| 16.07 | CTCATCGCCTCGTGGAATGGGC | Forward primer for qRT-PCR targeting the *metE* leader (upstream amplicon) | This study |
| 16.08 | TGCTGTCCTTCGATCGACGGGT | Reverse primer for qRT-PCR targeting the *metE* leader (upstream amplicon) | This study |
| 16.09 | AACCTTGTCTAGCGGTCGGGCC | Forward primer for qRT-PCR targeting the *ppe2* leader (upstream amplicon) | This study |
| 16.10 | CCATCCAGATCGGGGCGGTCAT | Reverse primer for qRT-PCR targeting the *metE* leader (upstream amplicon) | This study |
| 16.48 | ACTTGGATGTCGTGTTCGCT | Forward primer for PCR targeting the *cobQ-cobU* junction; used with 16.51 as reverse primer | This study |
| 16.49 | CTCAGCCAGGCTAGATCGG | Reverse primer for PCR targeting the *cobQ* coding region; used with 16.48 as forward primer | This study |
| 16.50 | GTGCCATCCCATTCTTCGGG | Forward primer for PCR targeting the *cobU* coding region, used with 16.51 as reverse primer | This study |
| 16.51 | TGACCAGATGTACCTCATCGC | Reverse primer for PCR targeting the *cobQ-cobU* junction | This study |

| **Gene Blocks Fragments** | | | |
| --- | --- | --- | --- |
| **Name** | **5’ → 3’ sequence** | **Description** | **Reference(s)** |
| gBlock TK-1 | ATTAGACTAAGCTTGCAGCTGGTCTGCTGGCGTCCGAAAGGGCGTCGGCATCGAGCGGGGCAACGATGCTTCGCGAGAGGGAACCTGGTGAGAATCCGGGACTGTCCCGCAGCGGTATGCAGGAACGACCGCCGTCTTGGAAGTAGACAAGCACTGGTCTCAACGACTGGGAAGCGACGGCCAGTAGGAGCACCCACCGGGTGCGAGCCTGCGAGTCCGAAGACCTGCCAGCCGTGCCGGACGCGCCGCGCCCGGCGGCTCATCGCCTCGTGGAATGGGCGTTTGGCCGTGCCTGTTGCCGGGTGCATCGGTGTGCACCTCAATCGGATCGGCTGCGCGTCCGCGGGCGGTGAACCACCCGTCGATCGAAGGACAGCACGTGACCATGGATGATCCCGTCGTTTTA | GeneBlocks fragment containing the full-length *metE* leader and the first two codons of MetE, used to generate pTKSW-*metE* by restriction cloning. *Hin*dIII and *Nco*I restriction sites are underlined. | This study |
| gBlock TK-2 | ATTAGACTAAGCTTGTCAGGCGATGACGATGCAGGAAGCCGGTGAGAATCCGGCGCGGTCCCGCCACTGTCACCGGGGAGCGACCCTCGTAAGCCACGGCCACAGGCTGGAAGGCGAGGCAAGCAACGATCCGGGAGCCAGGAGACTCGCGTCATCGCGTCCTGCCACCCGGGGCGGTGTACCCCGAGAAAGCTAACCGGTCATGACGCTCCAAACCTTGTCTAGCGGTCGGGCCACCACCACGCTGCTGGGCCTTAACGTGGCTGCTCGCGACGCGACGACACCTCCGCCGGCGGTCAGCCGCCGATGACCGCCCCGATCTGGATGGCTTCGCCCCCAGAGGTGCCCATGGATGATCCCGTCGTTTTA | GeneBlocks fragment containing the full-length *ppe2* leader sequence and the first 13 codons of PPE2, used to generate pTKSW-*ppe2* by restriction cloning. *Hin*dIII and *Nco*I restriction sites are underlined | This study |
| gBlock TK-3 | CCATTGCCGGATTTGTATTAGACTAAGCTTGCCGGGTGCATCGGTGTGCACCTCAATCGGATCGGCTGCGCGTCCGCGGGCGGTGAACCACCCGTCGATCGAAGGACAGCACGTGACCATGGATGATCCCGTTTTACAACGTC | GeneBlocks fragment containing the partial *metE* leader sequence, used to generate the SD-αSD construct by Gibson assembly. *Hin*dIII and *Nco*I restriction sites are underlined | This study |
| gBlock TK-4 | CCATTGCCGGATTTGTATTAGACTAAGCTTGCCCGGCGGCTCATCGCCTCGTGGAATGGGCGTTTGGCCGTGCCTGTTGCCGGGTGCATCGGTGTGCACCTCAATCGGATCGGCTGCGCGTCCGCGGGCGGTGAACCACCCGTCGATCGAAGGACAGCACGTGACCATGGATGATCCCGTTTTACAACGTC | GeneBlocks fragment containing the partial *metE* leader sequence, used to generate the SD-αSD-ααSD construct by Gibson assembly. *Hin*dIII and *Nco*I restriction sites are underlined | This study |
| gBlock TK-5 | CCATTGCCGGATTTGTATTAGACTAAGCTTGTCCCGCAGCGGTATGCAGGAACGACCGCCGTCTTGGAAGTAGACAAGCACTGGTCTCAACGACTGGGAAGCGACGGCCAGTAGGAGCACCCACCGGGTGCGAGCCTGCGAGTCCGAAGACCTGCCAGCCGTGCCGGACGCGCCGCGCCCGGCGGCTCATCGCCTCGTGGAATGGGCGTTTGGCCGTGCCTGTTGCCGGGTGCATCGGTGTGCACCTCAATCGGATCGGCTGCGCGTCCGCGGGCGGTGAACCACCCGTCGATCGAAGGACAGCACGTGACCATGGATGATCCCGTTTTACAACGTC | GeneBlocks fragment containing the partial *metE* leader sequence, used to generate the extended SD-αSD-ααSD construct by Gibson assembly. *Hin*dIII and *Nco*I restriction sites are underlined | This study |
